# Impact Induces Phagocytic Defect in Reactive Microglia

**DOI:** 10.1101/2025.09.14.676171

**Authors:** Ruilin Yu, Edmond A. Rogers, Palak Manchanda, Connor H. Beveridge, Caitlin E. Randolph, Timothy B. Beauclair, Krupal P. Jethava, Riyi Shi, Gaurav Chopra

**Author notes:** Corresponding author –. These authors contributed equally to the paper (Ruilin Yu, Edmond A. Rogers).

## Abstract

We have developed traumatic brain injury (TBI)-on-a-chip in vitro models using primary microglia and neuronal networks and recorded the molecular and cellular changes following impact to represent impact injury. Using a pH-responsive amyloid β (Aβ^pH^), we showed that microglial phagocytosis was reduced at 7 days post-impact on the chip. Simultaneously, neurons increased their uptake of Aβ, and decreased neuronal firing frequency at 7 days post-impact based on electrophysiological recordings. Given the importance of lipid metabolism in brain trauma and neurodegeneration, the lipidome secreted by impacted cells was analyzed to understand changes in cellular processes. Interestingly, many lipid species from the sphingomyelin, glycerophospholipid, and phosphatidylserine classes were significantly affected by impact, which are known to play important roles in the resolution of neuroinflammation and the pathogenesis of neurodegeneration.

## Introduction

Traumatic brain injury (TBI) is a leading cause of mortality, morbidity, and disability worldwide,^1,2^ and has been shown to significantly increase the lifetime risk of multiple neurodegenerative diseases, including Alzheimer’s Disease (AD). However, despite significant research efforts, there is currently no effective therapy to treat TBI or prevent the associated long term cognitive deficits, and the underlying mechanisms are still incompletely understood^3,4^. It has become evident that novel approaches will be required to uncover this complex pathogenesis, and while animal models have been useful to study TBI, their size, anatomical, and histological differences from humans have produced limited success with translation in clinical trials^5–7^. Further, while *in vitro* models typically offer the type of investigative resolution required for mechanistic studies, they historically lack the ability to withstand physical, traumatic forces to model TBI. In response, we recently developed an *in vitro* model of trauma termed “TBI-on-a-chip", capable of sustaining cells through a range of clinically relevant g-forces while offering morphological and electrophysiological monitoring for mechanistic studies and providing a high-throughput method for therapeutic testing^8–10^.

TBI can be divided into a primary, mechanical injury, and a subsequent secondary, biochemical injury. Recent studies suggest that some of these secondary injuries are capable of producing even more severe consequences than the original, primary insult, resulting in psychosocial problems, physical disabilities, and neurodegeneration^11–14^. While the initial, mechanical injuries are generally instantaneous (seconds) and relatively unavoidable, secondary injuries are long-lasting (hours to weeks), providing a unique opportunity for pharmaceutical or therapeutic interventions. Therefore, an improved understanding of TBI secondary injuries with *in vitro* models for drug screening could be vital to the generation of novel treatments. The TBI-on-a-chip system offers users the ability to isolate secondary biochemical changes from the mechanical impact, allowing for the investigation of media-tied biochemical changes (i.e. metabolite profiles) over chronic periods of 7 days post-injury. ^5–7^

Recent studies suggest that the brain’s lipidome could be the key to deciphering its complex structure and function^15^, a hypothesis that is strongly supported by the observation that lipids make up 50% of the brain’s dry mass^16,17^. Interestingly, TBI has been shown to disrupt the lipidome of plasma, cerebral spinal fluid, and brain tissue^18^. Further, due to its high levels of polyunsaturated fatty acids, the brain is particularly vulnerable to lipid peroxidation from reactive oxygen species. More specifically, acrolein, a demonstrated marker of central nervous system (CNS) trauma and suspected principal component of secondary injury^19,20^, is the most reactive α,β-unsaturated aldehyde with a variety of endogenous or environmental sources, including lipid peroxidation^21^. Furthermore, acrolein induces oxidative damage through rapidly reacting with proteins, lipids, DNA, or other biomolecules, which further propagates damage, creating a vicious cycle^19,22,23^. Despite the importance of lipids to brain health and homeostasis, to our knowledge no study has investigated the effects of physical injury on the lipidome of various classes of brain cells.

Microglia are the resident immune cells of the brain, comprising ∼10% of CNS cells in adults. They are essential for CNS immunosurveillance, clearance of waste, and injury repair, making microglia an emerging therapeutic target for various CNS pathologies^24–26^. Furthermore, microglia phagocytosis is essential to the resolution of neuroinflammation and neuronal survival in several models of CNS injury^27–29^. However, recent studies propose that injury-induced microglial phagocytosis may also lead to neuronal loss and memory deficit^30–32^. While increasing evidence suggests that lipid metabolism plays a critical role in driving the phenotypes and functions of microglia^33–35^, the underlying mechanisms and associated pathways have yet to be entirely elucidated. Several current investigations targeting lipid metabolism have shown that both ceramide and triacylglycerol synthesis are capable of significantly modulating microglial functions^36–40^. Therefore, improving our understanding of trauma, its effect on the lipidome of microglia, and the associated functional alterations, could provide significant insight into the pathogenesis of TBI, potentially revealing novel, more effective targets for intervention.

Here, we employed the morphological and electrophysiological capabilities of the TBI-on-a-chip system to reveal significant, concurrent deficits in both microglia and neuronal function at 7 days post-impact injury. We have integrated untargeted lipidome profiling to gain a better understanding of the molecular changes in physically injured cells. We found that disruption in sphingolipids and phosphatidylserine are correlated with impact-induced defects in the phagocytic ability of primary microglia and decreases in the firing frequency of primary neurons. Pathway analysis further revealed disturbances in glycerophospholipid, cholesterol, and sphingolipid metabolism pathways which may be involved in the injury-induced microglial and neuronal defects. We also noted significant increases in cellular (microglial and neuronal) acrolein-lysine adducts at 7 days post-injury. Exposure to a physiologically-relevant concentration of acrolein^20,41–43^ recapitulated the aforementioned impact-induced defects in microglia function and neuronal activity, suggesting that acrolein could play a crucial role in the secondary injury of TBI and the resulting neurodegenerative sequalae. Finally, it is our hope that by demonstrating a cellular platform capable of evaluating the function of multiple cell types, we can uncover unique insights into the underlying mechanisms that define impact-induced pathologies, individually and in concert, while creating an opportunity for further investigations into therapeutic opportunities.

## Results

### TBI-on-a-chip model optimized for histiotypical neuronal network and primary microglia

This study utilized “TBI-on-a-chip" for the application of all mechanical injuries. TBI-on-a-chip is a pendulum based, *in vitro* model of trauma, that is capable of generating consistent, rapid acceleration injuries through a clinically-relevant range of impact g-forces (30 – 300 g), while offering simultaneous electrophysiological and morphological monitoring (**Fig 1**)^8–10^. Two kinds of glass-based plates were used for the culture of histiotypical neuronal network and primary microglia. Multielectrode arrays (MEA) were used for culturing primary histiotypical cells and recording neuronal activity (**Fig 1AB, Fig S1A**). Enhanced Resolution Plates (ERP) were simplified plates which mirrored standard-MEA dimensions (5 x 5 cm) and strength to withstand physical impacts, while offering improved resolutions for morphological-focused investigations. Therefore, the ERPs were perfectly suited for culturing primary microglia (**Fig 1CD, Fig S1B**) or histiotypical cells for imaging purposes. For this investigation, we selected a moderate series of three rapidly administered 100 g impact injuries (4 – 6 second interval), to better focus on the complex interaction between repeated injury and the development of neurodegeneration^44,45^ (**Fig 1E, Fig S1C**). This impact force does not cause significant cell death or detachment, both shown previously and presently^9^. No significant difference in cell viability was found in impacted networks at 7 days post-impact (88.5 ± 0.9 viability, n = 5, p > 0.05) compared with sham networks (88.2 ± 1.0 %). Cell viability was quantified using fluorescein diacetate (FDA) and propidium iodide (PI) staining.

**Figure 1.**
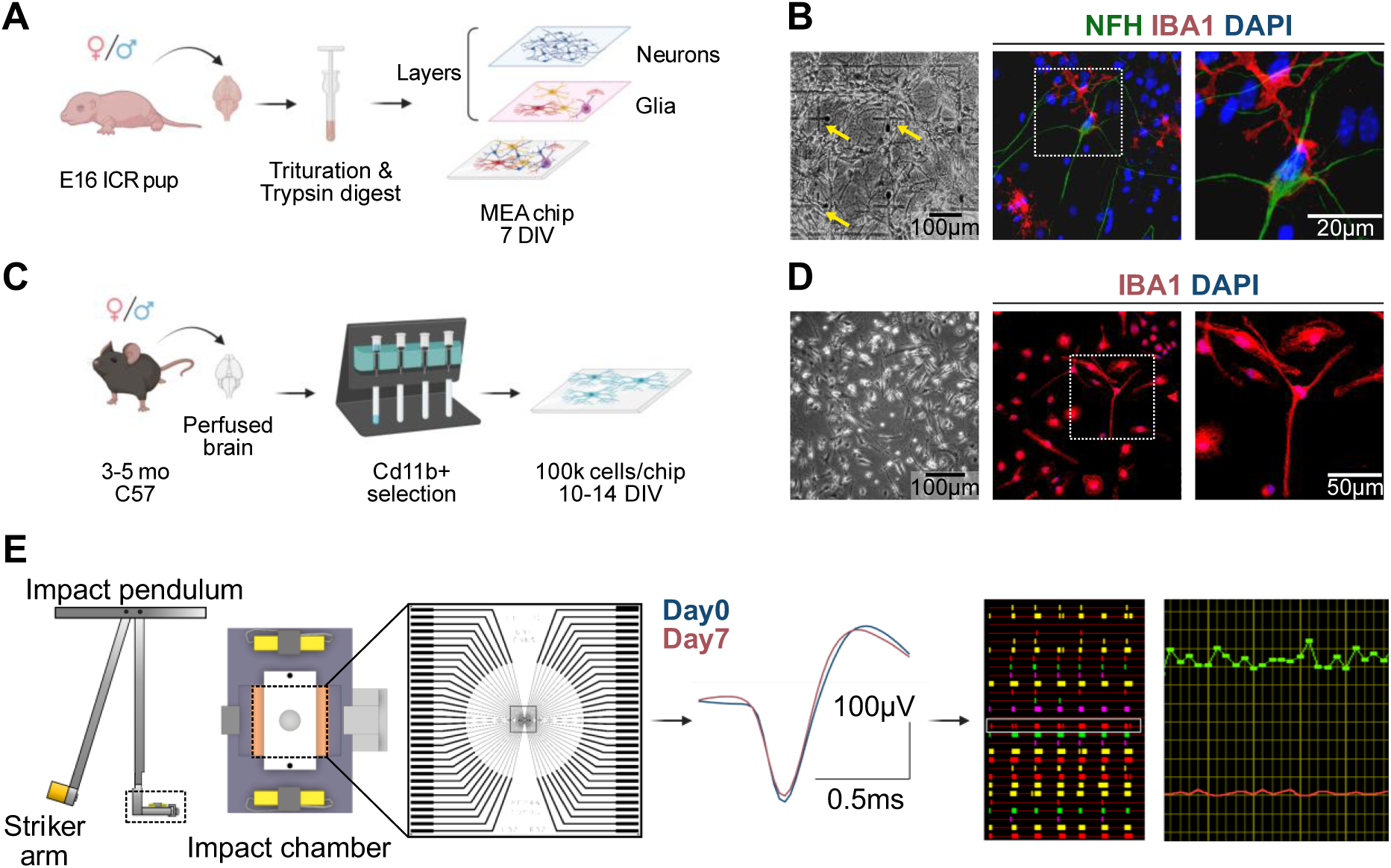
Overview of cell cultures and impact injury model. **(A)** Scheme of histiotypical neuronal network generation from E-16 murine embryos. **(B)** Phase-contrast images of neuronal network on multielectrode arrays (MEA). Yellow arrows indicate neurons growing over electrodes. Fluorescent microscopy images reveal co-culture composition: neurons (NFH, green), microglia (IBA1, red), and nuclei (DAPI, blue). **(C)** Scheme of primary microglia isolation from adult C57 mice via Cd11b+ magnetic-activated cell sorting. **(D)** Phase-contrast image of Cd11b+ selected microglia on enhanced resolution plates (ERP). Fluorescent microscopy images showed the morphology of homeostatic microglia (Iba1+) at 10-14 days *in vitro* (DIV). **(E)** Scheme of “TBI-on-a-chip” *in vitro* trauma model. The striker arm (SA) swings from a pre-selected height, contacting the impact-chamber/chip assembly, and generating a rapid acceleration injury of 100 g (3x). Sham-networks receive identical treatments, with the notable absence of the application of force. The impact chamber maintains physiological parameters during impact-injury while providing morphological and electrophysiological access to the network on MEA/ERP (square, dashed line, with typical MEA conductor pattern enlarged at right). Waveshapes from a representative unit pre (blue, Day 0) and 7 days post-impact (red, Day 7) show consistency, indicating stability of cell-electrode coupling during this 7 day period. This unit (white box) and others are visualized to the right via Plexon’s 2D raster plot display, which charts events (action potentials, y axis) against time (x axis). The corresponding spike frequencies are graphed as average spikes per minute (green line, ∼ 30 minutes shown) per total number of discriminated units (red line, 40 units), shown at the far right.

### Impact leads to reactive microglial phenotype with defective phagocytosis

Using *in vivo* models of traumatic brain injury (TBI), reactive microglia have been shown to adopt an amoeboid morphology and increase their release of nitric oxide (NO), a potent pro-inflammatory signal^46–48^. Consistent with these studies, impacted microglia in our model lost their processes and became amoeboid post-impact. Analysis of over 300 microglia confirmed that impact led to significant morphology changes at 7 days post-impact (DPI), characterized by reduced perimeter, morphological index (ratio of total cell area/soma area), compactness (roundness and area filled), and increased form factor (roundness) (**Fig 2B**). Additionally, increased NO levels were detected in the culture supernatant of impacted primary microglia at both 1 and 7 DPI (**Fig 2C, Fig S2B**).

**Figure 2.**
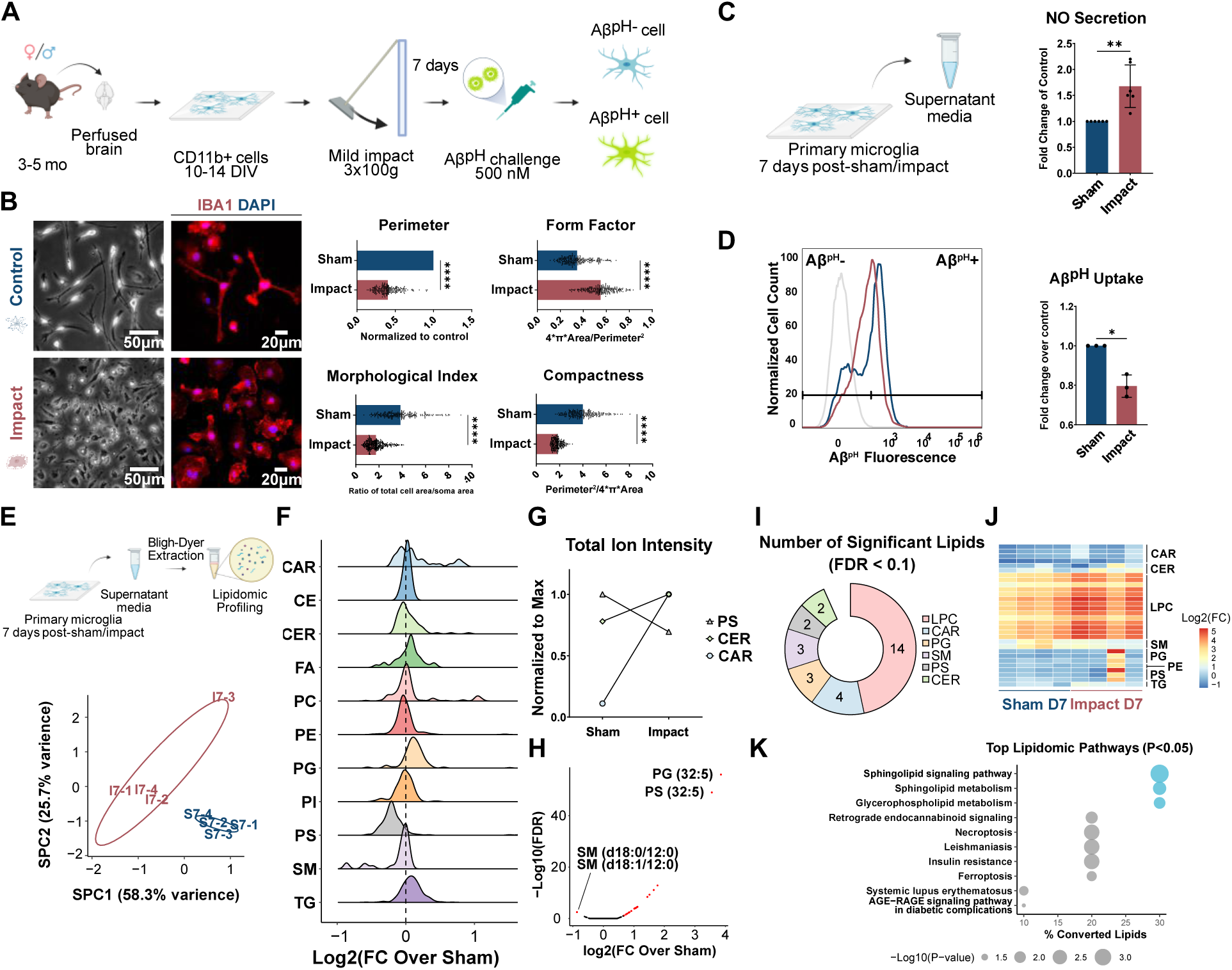
Impact induces phagocytic defect in primary microglia. **(A)** Scheme of primary microglia isolation, culture, impact treatment, and Aβ^pH^ challenge. **(B)** Quantified morphology of primary microglia 7 days post-sham/impact (DPS/I) from 20x phase-contrast microscopy. Around 300-400 microglia were profiled in total from each treatment group (n=3 biological replicates, mean±SE). Confocal images confirmed expression of microglia marker IBA1. **(C)** Nitric oxide (NO) levels increased in microglial supernatant 7 DPI (n=6 biological replicates, mean±SE). NO concentration in the supernatant media was determined using Griess assay and normalized to Sham. **(D)** Phagocytosis of Aβ^pH^ by impacted microglia on 7 DPI measured by flow cytometry. **(E)** Scheme of lipidomics profiling of microglial supernatant and Principal Component Analysis (PCA) analysis of 7 days post-Impact vs Sham (N=4 biological replicates). **(F)** Ridge plot of Log2-tranformed fold change (Impact/Sham) of all lipids from 11 major lipid classes. **(G)** Total ion intensity of ceramide (CER), sphingomyelin (SM), and acylcarnitine (CAR) normalized to respective maximum. **(H)** Volcano plot of significant lipids (red, FDR < 0.1) and their Log2-tranformed fold change in media from Impacted vs Sham microglia. Lipids with prominent fold changes are annotated. **(I)** Breakdown of significantly changed lipid species by class. **(J)** Heatmap of significantly changed lipid species. Color intensity corresponds to Log2(fold change). **(K)** Lipid classes with more than 2 significantly changed lipids were queued with *Mus Musculus* as background organism for LIpid Pathway Enrichment Analysis (LIPEA) pathway analysis^149^. Pathway results were ranked by percentage of converted lipids (number of queued lipids found in the pathway divided by all queued lipids). All error bars indicate SEM. Statistical significance was calculated with Student’s t-test, except for lipidomics data which was calculated with linear regression using the EdgeR package.

Microglial phagocytosis is a tightly-regulated and cargo-specific process^49,50^. We have previously developed a pH-responsive human Aβ1–42 analogue (Aβ^pH^) which detects cellular phagocytosis by fluorescence in the acidic environment of phagolysosome^51^. Several studies have found increased microglial phagocytosis of neuronal debris after acute injury^27,28^, which is consistent with their role as professional phagocytes in the CNS. However, we were surprised to find a significant decrease in microglial phagocytosis of Aβ^pH^ at 7 DPI, as shown through both flow cytometry (**Fig 2D**) and immunofluorescence imaging (**Fig 3D**). Previous studies have shown similar phagocytic impairment in reactive microglia in mouse models of Alzheimer’s Disease and Parkinson’s Disease^52,53^. Therefore, it is possible that inflammatory molecules induced by the impact generated the phagocytically impaired reactive microglia, similar to what was observed in the context of chronic neuroinflammation.

**Figure 3.**
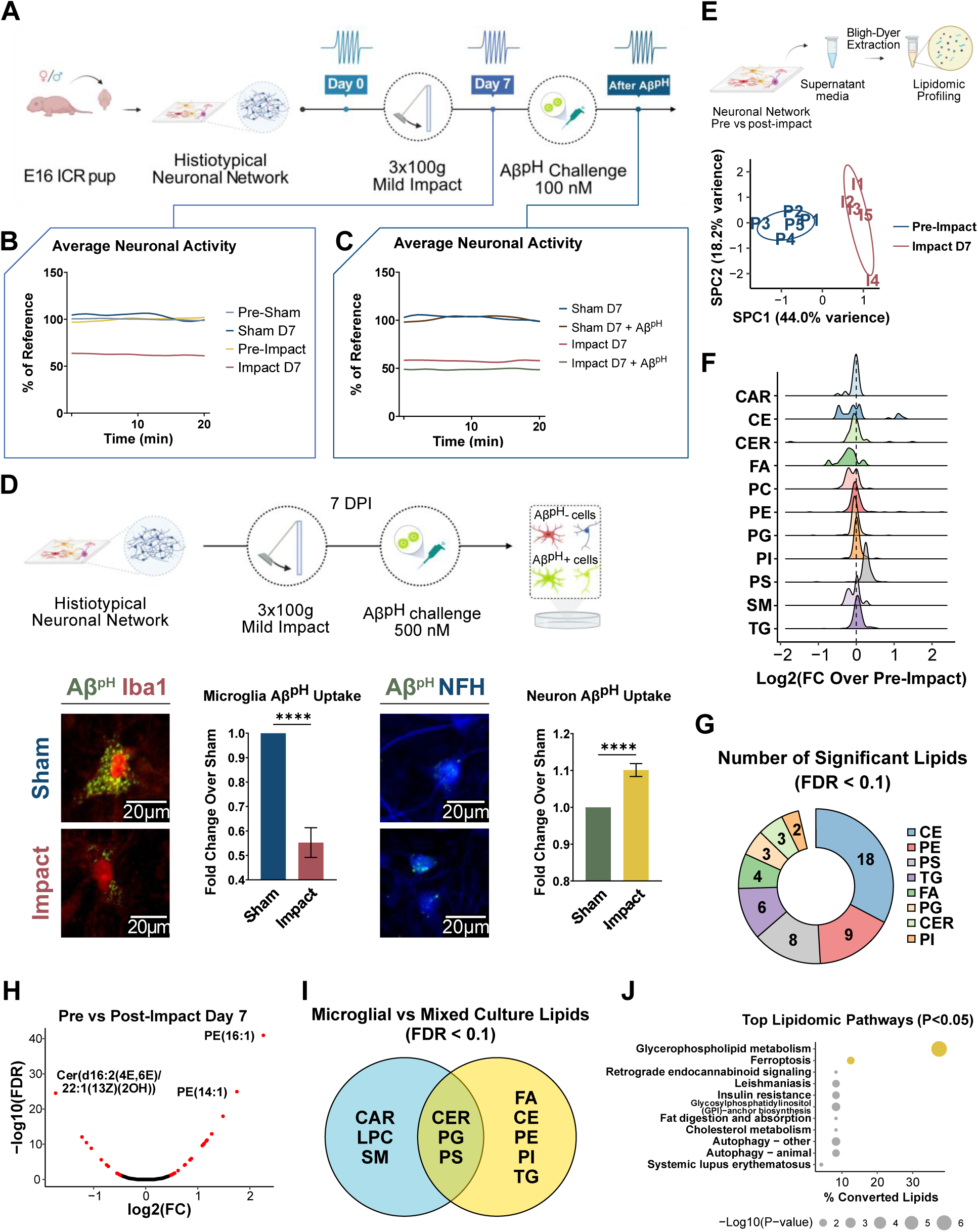
Impact induced defects in neuronal network activity and cell-type specific changes of Aβ^pH^ clearance. **(A)** Scheme of generation, impact, electrophysiological recording, and Aβ^pH^ challenge of neuronal networks at 7 days post-impact. **(B and C)** Electrophysiological recordings for consecutive 20-minute periods were collected on Day 0 (pre-impact), 7 days post-Impact, and 1 hour post-Aβ^pH^ challenge (n = 3 networks). Average network activity (spikes per minute, SPM) was normalized to network specific pre-treatment levels. **(D)** Aβ^pH^ (green) internalization by neurons (NFH, blue) and microglia (Iba1, red) in histiotypical networks 7 days after sham or impact (n=20). **(E)** Scheme of lipidomics profiling of mixed culture supernatant and Principal Component Analysis (PCA) analysis of Pre-Impact vs 7 days Post-Impact (n=5 networks). **(F)** Ridge plot of Log2-tranformed fold change (Impact/Pre-Impact) of all lipids from 11 major lipid classes. **(G)** Composition of significantly changed lipids (FDR < 0.1). **(H)** Volcano plot of significant lipids (red, FDR < 0.1) in Pre-Impact vs 7 Days Post-Impact samples. Lipids with prominent fold changes are annotated. **(I)** Venn diagram comparison of significantly changed lipids (FDR < 0.1) in primary microglia vs mixed cell cultures at 7 days post-impact. **(J)** Lipidomic pathway analysis via LIPEA^150^. All error bars indicate SEM. Statistical significance was calculated with Student’s t-test, except for lipidomics data which was calculated with linear regression using the EdgeR package^151^.

### Hypophagic microglia show disrupted sphingolipid metabolism, increased release of acylcarnitine and lysophosphatidylcholine, and decreased release of phosphatidylserine

To understand the state of hypophagic microglia at 7 DPI, we performed unbiased lipidomics on the supernatant media from sham and impacted microglia. Specifically, we detected 1,501 lipid from 11 classes in 200 µL supernatant culture media^40,54,55^. Principal component analysis (PCA) showed a clear distinction between the supernatant collected from microglia at 7 days post-Sham or Impact (**Fig 2E**), while the lipidome at 1-Day post-Sham or Impact considerably overlapped (**Fig S2E**), indicating time-dependent, progressive changes in microglia reactivity. The fold change distribution of acylcarnitine (CAR), ceramide (CER), phosphatidylglycerol (PG), and triacylglycerol (TG) species increased, while phosphatidylserine (PS) and sphingomyelin (SM) decreased (**Fig 2F**). Total ion intensity of CER and CAR increased, while all other lipid classes decreased, including PS (**Fig 2G**). A few species of PS and PG were among the most upregulated lipids, as indicated in the volcano plot (**Fig 2H**). Linear regression analysis revealed significant changes in many lipids at 7 days post-impact, highlighted by species of lysophosphatidylcholine (LPC), TG, CER, and SM (**Fig 2I**). Changes in lipid species from these classes accounted for 63.4% of all significant changes in the lipidome of impacted microglia supernatant (FDR<0.1). LIPEA pathway analysis indicated changes in SM signaling and metabolism, as well as glycerophospholipid metabolism. BIOPAN pathway analysis found significantly suppressed lipid conversion from PE to PS species, which could explain the overall decrease in PS (**Fig 2FG, Table S10**).

### Impact induced activity deficits and heightened Aβ sensitivity in neuronal networks

Impact injuries were applied to histiotypical neuronal networks on MEAs to investigate neuronal responses to trauma. Utilizing the electrophysiological capabilities of the TBI-on-a-chip system, native, pre-treatment network activity was established (Day 0) before “Sham” or “Impact” (3 x 100g) application, with subsequent analysis at Day 7 (**Fig 3A**). Consecutively sampled 20-minute segments of network activity (average spikes per minute) from Sham networks showed no significant deficits between Day 0 and Day 7 post-sham (p > 0.05, n = 3 networks, 51 total neurons) (**Fig 3B**). Interestingly, a minor increase in activity was observed. Identical evaluations of impacted networks revealed a significant reduction in average spike frequencies of 37.7 ± 9.9 % at Day 7 (p < 0.01, n = 3 networks, 60 total neurons, when compared to Impact Day 0 pre-treatment activity). Further, Day 7 recording experiments included the addition of Aβ^pH^ to culture media at 100 µM concentration, for determining any associated alterations of susceptibility for neuronal cells. Interestingly, Day 7 Impact networks showed additional, significant average deficits of 11.8 ± 4.1 % post-Aβ^pH^ addition compared to Impact Day 7 pre-Aβ^pH^ activity, while similarly exposed Sham cultures displayed no significant difference (**Fig 3C**).

### Impact reduced microglial and increased neuronal uptake of Aβ^pH^ in primary histiotypical networks

We performed immunofluorescent analyses of Impact and Sham histiotypical neuronal networks 7 days post-impact (3 x 100g) to determine cell-type specific phagocytosis of Aβ^pH^ **(Fig 3D)**. Consistent with flow cytometry results, quantification of relative light intensity measurements revealed an average reduction of 47.1 ± 5.94 % Aβ^pH^ uptake in impacted microglia when compared with procedurally and age-matched sham networks (p < 0.01, n = 20). In contrast, impacted neurons show average increases of 6 ± 4.45 % in Aβ^pH^ uptake, relative to controls (p < 0.01, n = 20). Qualitatively, the majority Aβ^pH^ uptake in Sham microglia was dense and soma-localized, with some appearance in more proximal segments of processes; impacts resulted in a marked, global-reduction of Aβ^pH^ signal from within microglia (Iba1). In Sham neurons, the appearance of Aβ^pH^ in was relatively sparse and multifocal, with subtle yet significant increase after Impact injury.

### Defective neuronal networks show disrupted sphingolipid, glycerophospholipid, and cholesterol metabolism

We next performed untargeted lipidomics on supernatant from impacted mixed histiotypical cultures to understand molecular changes that correlate with neuronal defects using supernatants isolated immediately before impact and at different times after impact. For all time points post-impact (4 hours, 3 days, and 7 days), the lipidome variation from impacted histiotypical neuronal networks was very different from pre-impact neuronal networks (**Fig 3E, Fig S3DH**). Interestingly, while Aβ^pH^ treatment on Day 7 post-impact did not fully separate the variation in lipidome, the Aβ^pH^-treated cultures had much less variance and clustered closely together compared to pre-treatment (**Fig S3L**) due to similar variation across all lipids. At 7 days post-impact (DPI) when microglial phagocytosis and neuronal activity reduction were observed, impact markedly increased the levels of cholesterol (CE), phosphatidylserine (PS), triacylglycerol (TG), and decreased the levels of free fatty acids (FA) released by mixed cells into the media (**Fig 3F**, **Fig S3G**). Changes in these lipid classes (CE, PS, TG, FA) made up 68% of all significantly changed lipid species (FDR<0.1, **Fig 3G**). A few species of PE were the most upregulated lipids at 7 days-post impact, and one species of ceramide was markedly decreased (**Fig 3H**). Interestingly, the composition of significantly changed lipids varied across different time points, with cholesterol esters present at all three time points (**Fig 3G, Fig S3FJN**). Many species of cholesterol esters (CE) were significantly affected at 7 days post-impact in mixed cultures (33% of all significant lipid species, **Fig 3G**), while they were not significantly affected in impacted primary microglia cultures. This is consistent with previous work that cholesterol synthesis in the CNS occurs primarily in neurons, astrocytes, and oligodendrocytes under homeostasis^56,57^. Overall, impact induced more diverse lipid changes in mixed culture (53 significant species from 8 classes) than in primary microglia culture (28 significant species from 6 classes), which coincides with more diversity in the cell types. Interestingly, the addition of Aβ^pH^ at 7 days post-impact in mixed cultures reversed the trend in the total ion intensity of FA, PE, CER, and PG classes (**Fig S3H**), implying that these lipids might play unique roles in Aβ^pH^-induced secondary injury.

Given the significant role of microglia in CNS injury response^46,48^, we expected overlap between the significantly changed lipid species from mixed histiotypical cells and primary microglia. While there were few overlaps in exact lipid species (**Table S3&8**), many major lipid classes were affected by impact in both types of primary cell cultures, including CER, PS, and PG (**Fig 3IJ**). Pathway analysis in mixed culture again revealed changes in glycerophospholipid metabolism, indicating that these pathways could be mediated by reactive microglia in the CNS context (**Fig 3J**). Additionally, changes in ferroptosis and autophagy pathways were found in mixed cultures.

### Acrolein impairs microglial phagocytosis and neuronal activity

Immunocytochemical studies performed on histiotypical neuronal networks experiencing either Impact (3x100g) or Sham treatments revealed significant increases in intracellular acrolein levels at Day 7 post-injury (compared to procedurally and age matched control networks), which was visualized utilizing fluorescent imaging (**Fig 4AB**). In summary, morphological analyses revealed the most pronounced elevations of acrolein in predominately perinuclear regions of both neuron and non-neuronal (glial) cells. While occasional observations in cellular processes were noted, it was at a relatively lower frequency. The results were quantified via immunostaining intensities and demonstrated a significant average increase of 11.9 ± 3.3 % (p < 0.01, n = 20, when compared to non-impact controls).

**Figure 4.**
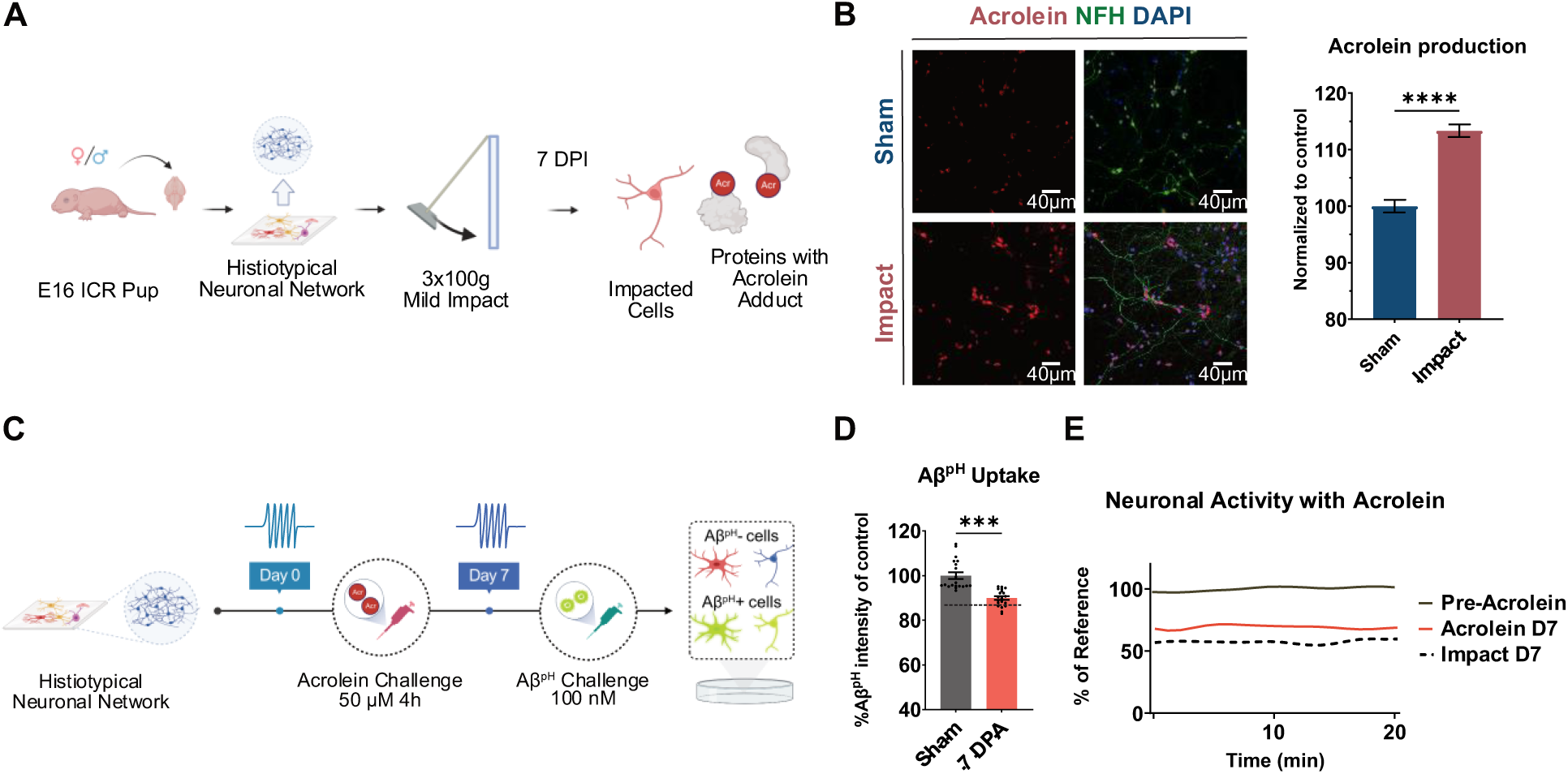
Acrolein is produced post-impact and impairs phagocytosis and neuronal activity. **(A)** Scheme of network generation, impact, and staining for acrolein-lysine adducts in histiotypical neuronal networks. **(B)** Representative immunofluorescent images of endogenously produced acrolein-lysine adducts (Stressmarq, red) in neurons (NFH, green) and glia cells from histiotypical networks 7 days after Impact or Sham. **(C)** Scheme of exogenous acrolein treatment, Aβ^pH^ challenge, and electrophysiological recordings of histiotypical networks. **(D)** Exogenous acrolein addition significantly reduced microglia phagocytosis of Aβ at 7 days post-acrolein treatment (7 DPA) (p < 0.01, n = 20 frames from 5 neuronal networks, 10.8 ± 4.4 %). **(E)** Electrophysiological recordings showed decreases in neuronal activity, measured as average spikes per minute over 20-minute periods and normalized to network specific pre-treatment levels (n=3 separate neuronal networks with 49 total neurons recorded, mean±SE). For comparison, impact day 7 activity results from figure 3 were indicated as the dashed line. Statistical significance was calculated using one-way ANOVA with Šídák’s post hoc test.

To investigate the role of acrolein in impact-induced changes, exogenous acrolein was introduced into media of non-impact histiotypical neuronal networks with subsequent analyses of Aβ^pH^ uptake or average activity at 7 days post exposure (**Fig 4CD**). Networks exposed to a 50 µM acrolein concentration for 4-hours showed average reductions in Aβ^pH^ uptake of 10.8 ± 4.4 % at Day 7, when compared to non-impact procedurally and age matched controls (p < 0.01, n = 20). Further, concurrently performed electrophysiological evaluations of acrolein-exposed networks revealed significant deficits in average spike frequencies of 30.1 ± 7.5 % at Day 7 (p < 0.01, n = 3 networks, 49 total neurons, when compared to Impact Day 1 pre-treatment activity, **Fig 4E**).

## Discussion

In this study, we utilized the novel TBI-on-a chip *in vitro* trauma model to produce clinically relevant impact-injuries in primary microglia and histiotypical neuronal networks. We have demonstrated significant, impact-induced functional alterations in both neurons and microglia at 7 days post-injury. To the best of our knowledge, this is the longest *in vitro* observation period reported in this type of cellular assay. These post-impact changes include an overall decrease of action potential production in neuronal networks, with reduction of phagocytic capabilities in microglia, as demonstrated by reduced uptake of Aβ^pH^ molecule^51^. In addition, we also detected disruption in sphingolipid, phospholipid, acylcarnitine, and cholesterol metabolism (**Fig 5**). Interestingly, exposure to Aβ(M1-42), a known modulator of neuronal activity^58–60^, at a concentration which produced no effect in non-impacted Sham networks, resulted in significant action potential deficits when introduced to neural networks at 7 days post-impact. Furthermore, we showed that the impact-induced elevations of acrolein-Lys adducts, a marker of oxidative stress and CNS injury in networks at 24 hours post-impact^9,61–63^, remain significantly elevated in all cell types (neuron and glia) at 7-days post-injury *in vitro*, a time point previously demonstrated *in vivo*^23,64,65^. We also showed that acrolein exposure alone was able to partially reproduce these impact-induced deficits in neuronal activity and phagocytic capabilities of microglia at 7 days post-exposure.

**Figure 5.**
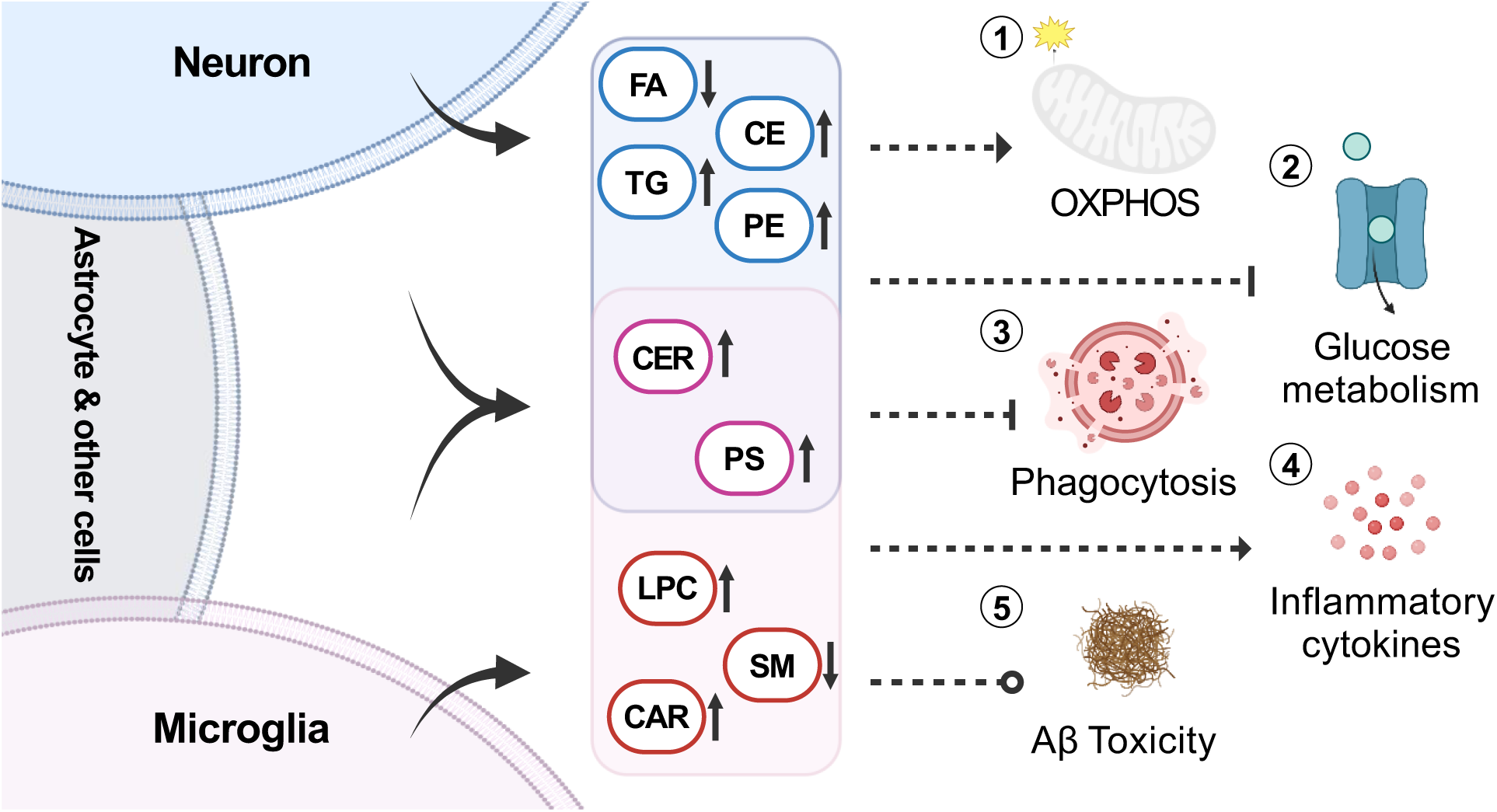
Proposed effects of impact-induced lipidomic changes on CNS cells. Impacted microglia release increased levels of acylcarnitine (CAR), lysophosphatidylcholine (LPC), and decreased levels of sphingomyelin (SM). Impacted neurons, astrocytes, and other cells from mixed culture released increased levels of cholesterol ester (CE), triacylglycerol (TG), phosphatidylethanolamine (PE), and decreased levels of free fatty acid (FA). Increased levels of ceramide (CER) were detected in both impacted microglia and mixed culture. Phosphatidylserine (PS) was increased in mixed culture but decreased in microglia post-impact. Potential effects of impact-induced lipid changes: 1) FA and TG fuel oxidative phosphorylation (OXPHOS) which is an alternative energy source and essential to phagocytosis in macrophages^89,90^. CAR is also neuroprotective by supporting OXPHOS, mitochondria functions, and normalizing cellular energy levels^101^. 2) FA, TG, and CER may inhibit glucose metabolism, leading to impaired neuronal activity^124,125^. 3) Extracellular CER and PS downregulate phagocytosis^83,92^. 4) CER, FA, and LPC can stimulate microglial release of pro-inflammatory cytokines^39,152,153^. Conversely, PS generally stimulates the release of anti-inflammatory cytokines^91^. 5) Cholesterol is neuroprotective while LPC exacerbates Aβ neurotoxicity^104,109^. Cholesterol is also important for stabilizing membrane structures for synaptic functions^108,154^.

Alterations in brain activities is one of the most sensitive measures of neuronal function. In the context of TBI, electrophysiological changes remain pronounced up to several weeks post-injury^66–68^. To detect the impact of injury in neuronal function, we have developed a modified protocol for the TBI-on-a-chip system capable of providing detailed readout of immediate^8^ and longer-term (7 day) electrophysiological and morphological alterations in histiotypical neuronal networks in response to clinically-relevant impact injuries. This expanded timeline for the TBI-on-a-chip model more closely aligns the system’s investigative capabilities with current, experimental timelines of primary and secondary injury, relative to comparative *in vivo* models^64,69,70^. It is our hope that this study will inspire more detailed investigations into subtle, TBI-induced cellular responses and activity changes, that will ultimately be scaled up to develop novel therapeutics for TBI intervention. The “TBI-on-a-chip” model may also be coupled with stem cell-derived human cell cultures to provide more clinically relevant findings.

After *in vitro* impact injury, we have observed a reactive microglia phenotype which is consistent with *in vivo* TBI studies, including the transition to amoeboid morphology, release of nitric oxide (NO), and changes in their phagocytic activity^27,30,47,48,71,72^. This confirms that microglia can detect and respond to physical impact and further implies that defective microglial clearance of Aβ could contribute to the secondary injury of TBI and potentially the development of neurodegenerative diseases. Effective phagocytosis requires numerous signaling pathways and structural changes for the successful recognition, internalization, maturation, and recycling of phagosomes and their cargos^72,73^, as defects in any of these steps can lead to impaired phagocytosis. Defective phagosome maturation/acidification is well documented in both microglia and neurons in AD^74–76^. A limitation in our phagocytosis assay is that Aβ^pH^ only reflects the amount of Aβ peptides taken into the effective phagolysosome (pH 4.5-5.0), whereas it cannot reflect which stages of phagocytosis or degradation were impaired^51^. Probes that specifically label the early phagosome (Rab5), late phagosome or lysosome (Lamp1, Rab7) can better indicate the rate and progression of phagocytosis^73^. Thus, we encourage future research to investigate which stage of microglial phagocytosis was impaired in the context of physical injury or neuroinflammation.

Abnormal lipid metabolism affects the fluidity of cell membranes and membrane-dependent processes such as phagocytosis and synaptic transmission, as well as a plethora of signaling pathways and overall cellular energy levels. We found that impacted microglia released increased levels of ceramide (CER), free fatty acid (FA), triacylglycerol (TG), and decreased levels of phosphatidylserine (PS) and sphingomyelin (SM) at 7 days post-impact. Differential expression analysis further revealed significant changes in species of lysophosphatidylcholine (LPC) and acylcarnitine (CAR). Disruption of sphingolipid metabolism is ubiquitous among a variety of neurodegenerative diseases and CNS trauma^77–79^. Accumulation of CER has been found in the brain from patients with AD^79^, as well as serum^80^ and brain^81,82^ from animal models of impact or blast TBI. Furthermore, extracellular CER has been shown to inhibit both endocytosis and the delivery of endocytosed material to lysosomes^83^. Both saturated and unsaturated FAs have been shown to increase microglial phagocytosis of Aβ1-42^84^, myelin^85^, and beads^86^ *in vitro*^87^. It is worth noting that these studies typically used a single species of FA with high concentration and purity, whereas we detected global increase in 36 species of FA, which could explain the different outcomes. Another study has shown that TG hydrolysis is required for maintaining cellular FA levels and efficient phagocytosis in macrophages, and that external supply of fatty acid or glucose could not normalize the phenotype^89^. In fact, alternative activation of macrophages (M2), characterized by enhanced phagocytic capacity, is known to rely on oxidative phosphorylation and fatty acid oxidation as their main source of energy, whereas classical activation (M1) is more dependent on aerobic glycolysis^90^. The same study also showed that TG uptake (through CD36 and lipoproteins) and lysosomal lipolysis is important for the formation and survival of M2 macrophages^90^. These studies led us to hypothesize that deficiency in TG uptake, lipolysis, fatty acid oxidation, and/or oxidative phosphorylation contributed to the impact-induced hypophagic microglial phenotype.

Interestingly, while impacted microglia released less PS, impacted mixed cells released more PS compared to sham. This is also reflected in BIOPAN analyses which predicted upregulation of phosphatidylserine decarboxylase (PISD, catalyzes the formation of PE from PS) and downregulation of PS synthases (PTDSS1 and PTDSS2) in the lipidome of impacted microglia, whereas opposite trends were predicted in the lipidome of impacted neuronal networks (**Table S10, Table S11**). Externalization of PS on the cell membrane is a well-known “eat-me” signal in apoptotic cells which stimulates phagocytosis and suppresses inflammation, resulting in “silent clearance” which is important for healthy homeostasis^91^. Addition of PS to the media prevented macrophage phagocytosis of red blood cells, potentially through competitive binding to PS receptors^92^. Although PS is not a consistent biomarker for CNS injury or diseases, PS delivery in liposomes or oral supplementation has shown various benefits for CNS injury and disorders^93,94^. Our results suggest that non-microglial cells may be releasing PS to dampen neuroinflammation and promote recovery, although increased PS release could inhibit microglial phagocytosis. However, it is worth noting that cells in the two culture systems were pre-exposed to signals which might modulate their response to impact injury, as they were derived from animals of different strain and age, and subsequently cultured in different media. While we observed decrease in microglial phagocytosis of Aβ in both systems, it is possible that different molecular pathways were involved.

Increase in CAR and LPC were unique to impacted microglia, as they were not significantly changed in impacted mixed culture. In a rat model of controlled cortical impact (CCI), mass spectrometry imaging has revealed increased levels of CAR in the brain, specifically colocalizing with the injury site where reactive microglia were found^95^. CAR is neuroprotective when administered after CCI^96^, ischemic injury^97^, or in the context of a variety of neurological disorders, including AD^98–100^. A suggested mechanism for the neuroprotective property of CAR is that it provides alternative fuel for neurons, which are in glucose shortage due to injury or disease. CAR also supports membrane synthesis and other cellular activities involved in injury repair^101^. Therefore, microglial release of CAR could serve neuroprotective function. Previous studies have shown that LPC were increased after repetitive, mild, cortical impact TBI^102^ and ischemia^103^. Interestingly, LPC increases the neurotoxicity and aggregation of Aβ1-42^104^ and is colocalized with various Aβ species in the AD mouse brain^105^. Overall, microglial lipidome is highly complex and likely plays multiple roles post-injury, regulating processes including phagocytosis, membrane repair, and fuel generation for themselves and other cells (**Fig 5**).

Lipids play important roles in modulating neurotransmission. Cholesteryl esters (CE) were the major lipid class in histiotypical cultures which were disrupted by impact. Cholesterol metabolism and transport has been strongly implicated in many neurodegenerative diseases, and the top risk allele for sporadic AD is a lipoprotein (ApoE) that transport CE between cells^106^. Recent study showed that chronic inhibition of cholesterol synthesis impaired neurotransmitter release probability^107^. Glial-derived cholesterol increases neuronal uptake of glutamate, an excitatory neurotransmitter^108^, and cholesterol enrichment protects neurons from the neurotoxicity of soluble Aβ^109^. It is generally believed that cells convert cholesterol into CE for either storage or transport^110,111^, and this conversion can be induced by lipid overloading and oxidative stress^112^. Accumulation of CE has been observed in excitotoxic brain injury^113^, AD^114^, and other CNS diseases^115^. Since we detected overall increase in CE from impacted mixed culture, it is possible that the cholesterol level was reduced in neuronal cell membrane which led to impaired synaptic functions.

Another possible explanation for defective neuronal firing activity is energy deficiency. Neurons rely heavily on efficient glucose uptake and metabolism for neuroactivity^116^. Disruption of glucose metabolism is well-documented in TBI patients, and the energy shortage can activate numerous pathways which contributes to AD pathogenesis^117,118^. Glucose transporter 1 (GLUT1) is lost after TBI in some studies^119,120^, although some others have found upregulation^117^. GLUT1 and GLUT3 expression were consistently reduced in post-mortem AD brains, and tracer studies have shown reduced glucose uptake in AD^121^. Pro-inflammatory stimuli, such as LPS or Aβ, increased microglial glucose uptake, which might exacerbate glucose shortage in the CNS parenchyma and impair neuronal health^122^. Neuronal glucose uptake and metabolism were also regulated by lipid signaling. For instance, CER can block the translocation of a glucose transporter (GLUT4) required for sustained neuronal firing^123,124^. In addition to CER, impacted mixed culture released increased levels of TG, phosphatidylethanolamine (PE) and decreased levels of FA. Elevated levels of TG and FA, often seen in diabetes patients, are long associated with decreased glucose consumption in muscle^125^, so they could contribute to glucose deficiency in the CNS. PE formation and oxidation is uniquely involved in ferroptosis^126^. Oxidized PE accumulated after cortical impact TBI in adult mice, and inhibition of 15-lipoxigenase was neuroprotective after TBI^127^. Interestingly, PE oxidation was also implicated in the phagocytosis of apoptotic cells and self-antigen presentation/tolerance in macrophages^128^, suggesting they may mediate injury resolution by microglia or astrocytes.

Finally, we have shown that acrolein plays an important role in secondary injury, due to its ability to partially recreate the impact induced deficits in neuronal activity, microglia phagocytosis, and even increased neuronal sensitivity to Aβ. As a demonstrated marker of CNS trauma and suspected component of AD-pathogenesis, acrolein has been reported by multiple studies to be increased after TBI, including using this model^9,10,64,129^. Acrolein has already been shown capable of altering both microglia and neuronal function^130–135^. Oxidative stress from other biochemicals such as 4-hydroxy-2-nonenal (4-HNE), another reactive aldehyde produced during lipid peroxidation, can impair the activity of lysosomal proteases^136^, or processes in the endocytosis^137^ or autophagy^138^ pathways which share many mechanisms with phagocytosis. Moreover, acrolein can promote ceramide production^139^, creating a complex network of inflammatory signals which propagate the damage. Pathway analysis of the lipidome from impacted histiotypical cultures also revealed ferroptosis and autophagy pathways. Ferroptosis is a form of cell death driven by excessive lipid peroxidation^140,141^, which contributes to neuronal death after TBI and in neurodegenerative diseases^127,142,143^. A previous study using a triculture system consisting of microglia, astrocyte, and neuron has shown that the presence of microglia exacerbated ferroptotic neuronal death^143^. However, microglia are capable of resisting ferroptosis through the activity of inducible nitric oxide synthase (iNOS) which inhibits phospholipid peroxidation^144^, but it is unknown whether they can “detox” surrounding neurons through this mechanism. Autophagy can regulate ferroptosis by regulating the available lipid pool through lipophagy and lipid recycling^145,146^. In controlled cortical impact TBI of young adult mice (9-12 weeks), microglia accumulated autophagy markers (LC2, SQSTM1), indicating inhibited autophagic flux^147^. When microglial autophagy was disrupted in AD, microglia lost their ability to proliferate and engage with Aβ plaques, as well as adopting senescent phenotypes^148^. Altogether, our results revealed a complex network of cellular processes which might contribute to the secondary injury after impact (**Fig 5**).

## Acknowledgement

We thank the following individuals for their valuable input and assistance: Ms. Anisa Dunham for help with animal breeding and colony management; Dr. J. Paul Robinson and Ms. Kathy Ragheb at the Purdue University Cytometry Laboratories for flow cytometry resources. Jhon Martinez and Shatha Mufti for their help in preparing electrophysiological experiments. This work was supported by the United States Department of Defense USAMRAA award W81XWH2010665 through the Peer Reviewed Alzheimer’s Research Program, and, in part, by the National Institutes of Health (NIH) award from National Center for Advancing Translational Sciences U18TR004146 award, and ASPIRE Challenge and Reduction-to-Practice awards to G.C. Additional support, in part, by the Department of Health, State of Indiana; the Indiana Clinical and Translational Sciences Institute grant UL1TR002529 is acknowledged. In addition, a portion of this electrophysiological research was made possible by a generous donation of equipment from the University of North Texas and Harvey Wiggins of Plexon, Inc. We also thank current and prior members of the Chopra and Shi laboratories for critical discussions and day-to-day assistance with experiments included in this manuscript. G.C. is the James Tarpo Jr. and Margaret Tarpo Professor of Chemistry.

## Author contribution

Conceptualization, R.Y., E.R., P.M. R.S., and G.C.; Methodology, R.Y., E.R., P.M., C.B., C.R., and T.B.; Software, R.Y., E.R., C.B., and C.R.; Formal Analysis, R.Y., E.R., and C.B.; Investigation, R.Y., E.R., P.M., and T.B.; Resources, R.Y., K.J., E.R., and T.B.; Data Curation, R.Y., E.R., C.B., and C.R.; Writing – Original Draft, R.Y. and E.R.; Writing – Review & Editing, R.Y., E.R., C.B., C.R., R.S., and G.C.; Visualization, R.Y., E.R., and C.B.; Supervision, R.S. and G.C.; Funding Acquisition, R.S. and G.C.; Project Administration, G.C.

## Declaration of interests

G.C. is the Director of Merck-Purdue Center funded by Merck & Co. and co-founder of Meditati Inc., Braingnosis Inc. and LIPOS BIO Inc. All other authors declare no competing interests.

## STAR★Methods Resource availability

### Lead contact

Further information and requests for resources should be directed to the lead contact, Gaurav Chopra

### Materials availability

*All materials used in the paper are commercially available or previously published. In-house generated materials, including usage of “TBI-on-a-chip” device, MEA/ERP chips, or Aβ^pH^ phagocytosis marker, can be made available from the lead contact upon request*.

### Data and code availability

- Data reported in this paper and any additional information required to reanalyze the data reported in this work are available upon request from the lead contact.

Analysis codes for the lipidomics and metabolomics experiments are available on Github at https://github.com/chopralab/Impact-Microglia-2024

## Method details

### Quantification and statistical analysis

Statistical analysis in this study was performed using softwares listed in the KRT and appropriate statistical tests chosen for each type of experiment. In general, t-test was used for comparison between two groups; one-way ANOVA with post hoc test was used for comparison between more than two groups. Statistical details can be found in the figure legends, results, and method section for each type of experiment. The value of definition of n, definition of center, and dispersion measures are described in the methods. Significance was defined based on calculated p-value and set threshold, which can be found in the figure legends. No data was excluded in the analysis unless indicated otherwise in the methods.

### Animal Ethics

Animal maintenance and isolation of primary cells (histiotypical and microglia-only) were performed according to Purdue Animal Care and Use Committee guidelines and approval (protocols #2003002027 and 1306999879), as well as the recommendations in the ARRIVE guidelines. C57BL/6J mice (Jackson Laboratory) and ICR mice (Envigo Inc.) were housed in pathogen free facilities with controlled temperature (22°C), humidity, lighting (12-h light/dark cycle), and food and water *ab libitum*.

### Microelectrode array preparation

Microelectrode arrays (MEA) were fabricated in-house utilizing standard photoetching techniques that have been described in previous publications^8–10^ and summarized in **Fig S1A**. Briefly, indium tin oxide (ITO) plates were photoetched, spin-insulated via polysiloxane, cured, and laser de-insulated at central electrode termini creating shallow (∼ 2 μm deep) recording craters, that were subsequently electrolytically gold-plated to lower the interface impedance to ∼ 1 MΩ. The hydrophobic insulation material (methyl-trimethoxysilane) resin insulation was mask-flamed for localized surface activation and sequentially coated with poly-D-lysine and laminin^155^. The conductor patterns used for MEAs consist of 64 semi-transparent ITO electrodes arranged in either a single or dual recording matrix configuration (**Fig 1**)^156^. The electrophysiological experiments performed in this study utilized both designs. In addition, this study also employed a novel, modified-MEA design, termed “Enhanced Resolution Plates”, or ERP. These unique, simplified plates were separated from T1X0015 (Colorado Concept Coatings, LLC), or soda lime float glass (non-polished) with a sputtered quartz barrier (1,000 Å), and mirror standard-MEA dimensions (5 x 5 cm), while producing an overall thinner substrate (∼1 mm) that is much more uniform than its insulated counterpart. The result is an MEA-substitute that is still capable of surviving high-impact forces, while offering moderately-improved resolutions for morphological-focused investigations. Further, these plates reduce the exposure of sensitive electrophysiological technologies with the relatively-harsh chemical treatments associated with some fixing/staining techniques.

### Primary histiotypical cultures

Frontal cortex tissues harvested from E-16 murine embryos of ICR mice (Envigo, Inc) were disassociated, pooled, and seeded onto MEAs or ERPs, using previously established protocols with minor modifications (**Fig 1A**)^8,157,158^. Briefly, cortices were isolated, mechanically separated, enzymatically digested with a 0.05 % trypsin solution (Gibco), triturated, combined with Dulbecco’s Modified Minimal Essential Medium (DMEM, HyClone), supplemented with 4 % fetal bovine serum (FBS, Gibco) and 5 % horse serum (Gibco), and seeded onto MEAs or ERPs at ∼150 k or 30 – 50 k cells per 100 µL, respectively. Networks were incubated at 37 ° C in a 10% CO2 atmosphere for 2 days post-seeding, and then transitioned to media without FBS, followed by biweekly media changes. The result is a histiotypical culture, providing a mixture of neuronal and glial cell types that are representative of the parent tissue (**Fig 1B**)^159^. All networks were allowed to mature 7 days before any treatment application, unless otherwise noted (see “Recording”).

### Primary microglia culture

Our primary microglia isolation and culture protocol was derived from Collins and Bohlen^160^ with minor modifications for adult mice. Briefly, 12-20 weeks old C57B/6 mice of either sex were euthanized with CO_2_ asphyxiation for 2.5-3 min at level 2.5-3 L/min. To eliminate macrophages in the brain, transcardial perfusion with cold PBS was performed until the liver completely discolors and the brain is milky white. Perfused brains were cut into 1 mm^3^ pieces before homogenization in Dulbecco’s Phosphate Buffered Saline++ (DPBS with Ca2+ and Mg2+ ions, Gibco) containing 0.4% DNase I (Worthington #LS006343) on the tissue dissociator at 37°C for 35 mins (Miltenyi Biotec). Cell suspension was filtered through a 70-μm filter. Myelin was removed in two rounds: first with gradient centrifugation with 23% Percoll (Cytiva) in DPBS (Gibco), then by myelin removal magnetic beads using LS columns (Miltenyi Biotec). After complete myelin removal, CD11b+ cells were positively selected from the single cell suspension using the CD11b beads as per the manufacturer’s instructions (Miltenyi Biotec). The CD11b+ cells were finally resuspended in 10% CO_2_-equilibrated microglia growth media (MGM) made in DMEM/F12, phenol-red free (Gibco #21-041-025).

Meanwhile, ERPs were sequentially coated with Poly-D-Lysine (10ug/mL in ultrapure water) for 1 hour at RT, triple washed with UP-H_2_O, and coated with collagen (2 ug/mL in MGM) for 1 hour at 37°C. The collagen was aspirated and 1x10^5^ cells were spot seeded in the center of each chip followed by incubation at 37°C for 10 minutes. The cells were topped with ∼1.5 mL TIC media (TGF-β, IL-34, cholesterol)^160^ supplemented with 2% fetal bovine serum (R&D Systems #S11195H). Primary microglia were maintained at 37°C and 10% CO_2_ with ∼50% media change every other day until the day of sham/impact treatment (around 10–14 div). Cells were maintained for 1 to 7 more days after sham/impact with ∼50% media change every other day. MGM and TIC media were prepared fresh monthly. The entire procedure of primary microglia isolation and culture on ERPs is summarized in **Fig S1B.**

### “TBI-on-a-chip” Injury Device

This study utilized “TBI-on-a-chip" for the application of all mechanical injuries. This approach necessitates a semi-portable life-support system capable of protecting the MEA/ERP adhered network during periods of high g-force exposures, which is accomplished using our stainless steel “miniature-incubator” impact-chamber (**Fig 1E**). Prior to impact, the MEA/ERP attached cells are transferred from standard incubation to the impact chamber, which is then secured to the target arm of the pendulum and impacted via the striker arm. While TBI-on-a-chip is capable of generating a range of clinically-relevant impact g-forces, for this study we employed 3 rapidly (4 – 6 sec) administered mild (100g) impacts. Following impact, the cells are removed from the impact chamber and returned to the incubator. In a non-electrophysiological based experiment, the entire procedure is completed in approximately 5 minutes. All experiments are performed in tandem with age and procedurally matched Shams^8-10^.

### Recording

All network electrophysiology was performed utilizing previously described techniques, with some minor modifications^8,10,156,161^. This method employs the use of neuronal networks (average age: 26 ± 5 days *in vitro*) cultured onto MEAs that are aseptically mounted inside proprietary stainless-steel recording chambers with concomitant life-support features. Taken together, this system provides consistent, precise command of the associated physiological parameters. This includes maintaining temperature (37 °C), pH (7.4, via continuous flow of 10% CO_2_ in air at a rate of 10 mL/min), and osmolarity (300 mOsmol/kg, using a constant infusion of ultrapure water).

A 64-channel, dual amplifier system was utilized to record spontaneous extracellular activity (Multichannel Acquisition Processor System, Plexon, Inc., Dallas, TX). Analog electrical signals were amplified (10 k total gain) and digitized (40 kHz), with time stamp conversion for storage and post-analysis (25 µs resolution). Real-time identification and sorting of active units was performed using Plexon waveshape templates. Under optimal conditions (visual signal-to-noise ratios > 3 : 1), this system can distinguish up to four separate waveforms from a single electrode in real-time. Network activity via spike production was quantified as previously described^8,157,162^. Briefly, spike production for each network was plotted as mean spikes per minute for the entire duration of the experiment. For each minute, the activity total was divided by the number of discriminated “active channels”. Active channels were defined as channels that reached at least 10 discriminated spike signals per minute, which is relatively conservative. Taken together, this method provides real-time monitoring of the evolution of network activity, while offering additional analysis capabilities offline.

Each network’s spontaneous activity is distinct; thus, data is expressed as percent change from network-specific, pre-treatment reference activity, which is maintained at its native, stable state for a minimum of 60 minutes before any experimental manipulation. Offline analysis of neurophysiological parameters was performed utilizing Plexon raster display, SortClient, and Neuroexplorer. (NEX Technologies, Colorado Springs).

To accomplish the 7 day recordings featured in this investigation, a slightly modified protocol was introduced. In summary, initial pre-treatment (Day 0) network recordings were performed as previously described. After the application of each respective treatment, networks were returned to standard-incubation conditions, where they continued to receive the standard care described under “Primary histiotypical cultures.” At 7 days post-treatment, networks were identically prepared and evaluated using matching recording parameters, before subsequent exposure to 100 nM of fluorophore-conjugated β-amyloid 42 (Aβ^pH^), a typically non-toxic concentration that is also the approximate threshold at which aggregation begins^51,163^. It is important to note, that minor changes can occur during this 7 day period, therefore user-matching of waveshape templates (Plexon) was paramount. This tedious approach is highly discriminative, and thus ultimately results in reduced total channel counts (average of 18 ± 3 channels, n = 9 networks). Every procedure was control-validated.

Network activities were quantified with Plexon/NeuroExplorer and were shown as % of pre-treatment reference activity values ± SD (n = 3 separate networks per treatment). Significance between treatments was established using an unpaired Student’s *t-*test, and normality was tested for using Shapiro-Wilk test. Statistical significance was defined for all as p value < 0.05. StataSE16, Microsoft Excel, and Graphpad Prism^164^ were used to perform statistical analysis.

### Preparation of Aβ^pH^

Recombinant human Aβ(M1-42) were produced in *E. coli* BL21(DE3) and purified according to our published protocol^165^. Competent *E. coli* BL21(DE3) (Invitrogen, Cat# C606010) were transformed with pET-Sac-Abeta(M1-42) plasmid (Addgene, Cat#71875). Successful transformants were selected and grown until OD=0.45, and the expression of Aβ(M1-42) was induced with 0.1 mM IPTG. After cell harvest and lysis, Aβ(M1-42) in inclusion bodies were solubilized in 8M urea buffer and purified with reverse-phase HPLC. Purified recombinant Aβ(M1-42) were conjugated to Protonex Green™ (PTX) as previously described^51^. The Aβ was monomerized in hexafluoroisopropanol (HFIP) and reconstituted in 1 M NaHCO3 (1 mg/mL, pH∼8.3). The Aβ were allowed to react with 10 molar equivalents of PTX (5 mM in anhydrous DMSO) for 3h in the dark at room temperature. Then, 5 mol eq. of PTX was added, and the reaction was incubated for 3 more hours under the same conditions. The conjugation product (Aβ^pH^) was washed and concentrated using Pierce Protein Concentrator columns (3K molecular weight cut off, ThermoFisher, Cat #PI88514) in ultra-pure water. MALFI-TOF MS was used to verify the m/z of the peptide product (4645 g/mol for Aβ(M1-42), 5070 g/mol for conjugated Aβ^pH^). To aggregate Aβ^pH^, the peptides were first dissolved in DMSO until 5 mM, then diluted in TIC media until 100 µM, immediately sterile filtered, and incubated in 37°C for 24 hours in the dark to aggregate. This solution was then aliquoted and stored in -80°C until usage.

### Immunofluorescence of histiotypical networks

All networks were thrice rinsed in phosphate buffered saline (PBS, pH = 7.4, Gibco) and fixed in 4% paraformaldehyde (PFA, Thermo Scientific) at 7 days post-treatment. Cultures were then washed with PBS, permeabilized with 0.2% Triton (Sigma), blocked in 10% normal donkey serum blocking solution (Abcam, AB7475), and treated overnight with primary antibodies at 4°C. Next, cells were gently rinsed with 0.1% Tween 20 (Sigma), treated with secondary antibodies for 2 hours, rinsed once more with tween, and bathed in PBS for imaging. This study utilized the following antibodies and stains: Iba1 (Wako, 019-19741); Alexa Fluor 594 (Jackson Immunoresearch, 711-585-152); Alexa Fluor 488 (Jackson Immunoresearch, 703-545-155); Anti-NFH (Neurofilament Heavy Chain) (Aves Labs, NFH); mouse anti-acrolein (Stressmarq, SMC-504D); and DAPI (Thermo Fisher, 62248). In addition, these experiments also used Aβ^pH^, a novel, custom-made human Aβ42 analogue, which exhibits green fluorescence after internalization of acidic organelles in cells but not at physiological pH^51^. Aβ^pH^-focused morphological experiments used a 1 hour, 500 nM exposure to better facilitate visual analysis, while still retaining a relatively innocuous total concentration and exposure period^51,58^. Quantitative measurements utilized the relative light intensity of Aβ^pH^ or acrolein-lysine adduct (acrolein-Lys) levels, using a customized Olympus IX81 fluorescence motorized phase contrast microscope with a Qimaging EXi Blue CCD camera and MetaMorph Premier. Image analysis was performed with ImageJ. Representative fluorescent images were taken with a Zeiss LSM 880 Microscope with Airyscan (Carl Zeiss, Germany) using ZEN black software.

Immunocytochemical and fluorescent data were analyzed as described previously^9^. Briefly, quantitative immunofluorescent data was obtained and averaged from 4 non-overlapping sections in 5 separate networks, or 20 total frames per treatment. Relative light intensity measurements were quantified with ImageJ (NIH) and are shown as % control values ± SD (n = 20 frames). All data was exported into GraphPad Prism^164^ for visualization and analysis. Statistical significance was calculated using unpaired t-test or one-way ANOVA with Šídák’s post hoc test.

### Immunofluorescence of primary microglia

0.01% Triton in PBS was used for all washes unless specified. Cells were permeabilize with 2% Triton for 30 min, washed 3x, and blocked with 1% normal goat serum (NGS) in PBS for 30 min. Cells were incubated with primary antibody (Wako Iba1 rabbit-anti-mouse, 1:500 in 1% NGS) overnight at 4°C. The next day, cells were 3x washed and incubated with secondary antibody (Invitrogen goat-anti-rabbit Alexa fluor 594, 1:1000 in 1% NGS) for 1 h at RT. After Iba1 staining, cells were washed 3x and incubated with 300 nM DAPI (1:1000 in PBS from 300 uM stock in water) for 5 min at RT. Finally, cells were washed 3x in PBS and mounted on Fluoromount G Anti-Fade (Southern Biotech #0100-35) and cover slides. Mounted cells were dried overnight at RT. The next day, 20-30 z-stack images were taken on Zeiss880 confocal microscope with step-width 0.2 μm using ZEN black software. The images were taken using 20x, 40x water immersion lens, or 63x oil immersion lens with various digital zooms. Z-stack (.czi) files were processed consistently on ImageJ FIJI^166^.

### Morphology analysis of primary microglia

Ten microscope images were taken on phase-contrast microscope with 20x lens at 7 days post-impact or sham. Between 200-300 cells were selected and analyzed in these images using Cell Profiler from n=3 biological replicates in independent experiments^167^. Specifically, cell soma was selected as “Primary Objects”, and entire cells including processes were selected as “Secondary Objects”. Then, the auto-selected objects were manually corrected. Corrected secondary objects were used to calculate the following features: perimeter, form factor, and compactness. Morphological index was calculated as total cell area/cell soma area, and it equals 1 for a cell with no cytoplasm. Form factor was calculated as 4*π*Area/Perimeter2, and it equals 1 for a perfect, filled circle. Compactness was calculated as the mean squared distance of the object’s pixels from the centroid divided by the area; compactness equals to 1 for a perfect, filled circle, and it is greater than 1 for irregular objects or objects with holes. All data was exported into GraphPad Prism^164^ for visualization and analysis. Statistical significance was calculated using unpaired t-test.

### Flow cytometry analysis of primary microglia

For Aβ^pH^ uptake, primary microglia were treated with 500 nM of Aβ^pH^ aggregates in TIC media in a 37°C incubator for 1 hour. After staining, the cells were washed once with room temperature PBS and detached in ice-cold PBS by repeated pipetting. Immediately before flow cytometry analysis, cells were stained with DAPI (0.1 μg/mL cell suspension) for 3 min on ice. Cells were analyzed on the Attune NxT flow cytometer (Invitrogen). The files were then analyzed on FlowJo V10 software^168^. All data was exported into GraphPad Prism for visualization and analysis. Statistical significance was calculated using unpaired t-test.

### Exogenous Acrolein Exposure

Upon maturation, networks were exposed to a 50 µM concentration of acrolein (Sigma) for a period of 4 hours, with procedurally and age matched controls. As described previously, a 100 mM stock solution was prepared for each experiment, and diluted in culture media to the final, physiologically-relevant concentration of 50 uM^20,41–43^. Acrolein is highly reactive, and thus stock solutions were prepared fresh for each experimental series (never stored)^20,169^. After the 4-hour exposure period, media was replaced with standard culture media and returned to incubation. At 7 days post-exposure, networks were electrophysiologically evaluated or exposed to Aβ^pH^ (500 nM for 1-hour), before subsequent rinsing (3 x PBS), fixation (4%PFA), and morphological analysis. All culture imaging was performed in PBS.

### Lipidomics

Supernatant was collected from cells and stored at -80C. Once all biological replicates were acquired, lipid was extracted from the samples using the Bligh-Dryer extraction according to previously established protocols^40,54,55^. Briefly, supernatant was mixed with methanol and HPLC grade chloroform (2:4.5:2.5) and incubated at 4℃ for 15 min. Then, equal volume of water and chloroform (2.5:2.5) was added to the mixture, resulting in a biphasic solution. Samples were centrifuged at 16,000xG for 10 min. The resulting mixture should consist of three layers: top layer containing lipids, interphase containing protein, and bottom layer containing metabolites. After each sample component was collected, the solvent was evaporated using speed-vac, leaving the dry lipid or metabolite extract in the bottom. Dried lipid extract was stored in -80C until analysis. Multiple reaction monitoring profiling of the extracted lipids was performed as described previously^40,54,55^. To prepare the lipid samples for MS injection, lipid extracts were reconstituted in 200 μl methanol:chloroform (3:1 v/v) and further diluted 100x in injection solvent (7:3 methanol:acetonitrile with 10 mM ammonium formate) in glass LC vials with inserts. The injection solvent alone without any lipids or metabolites was used as the ‘blank’ sample. MS data were acquired by flow injection (no chromatographic separation) from 8 μl of diluted lipid extract stock solution delivered (per sample per method) to the jet stream technology ion source (AJS) of an Agilent 6495C Triple Quadrupole mass spectrometer. This method enabled the interrogation of the relative amounts of numerous lipid species within 11 major lipid classes based on the LipidMaps database. See Supplementary Table 1 and 2 for total number and identity of lipids screened. EdgeR package was used to perform statistical analysis^151^. This method has been described in a previous study^54^. EdgeR uses a generalized linear model (GLM) to identify differentially expressed lipids. The GLM is based on the negative binomial distribution and allows the model to handle the technical and biological variability with a dispersion term using the common dispersion method^170^. Lipids were considered based on a false discovery rate (FDR) value <0.1^171^.

### Griess assay for NO quantification

The nitric oxide (NO) production was quantified colorimetrically using Griess assay. The supernatant from cells were collected, vortexed, and triplicate of 50 µL was aliquoted in a 96-well plate. A standard curve is prepared using sodium nitrate (0-80 µM NaNO_2_). 25 µL of Griess Reagent A (1% sulfanilamide in 5% H_3_PO_4_/water) was added to each well. Then, after 5 min of incubation on ice, 25 µL of Griess Reagent B (0.1% N-(1-Naphthyl)-ethylenediamine dihydrochloride (NED) in water) was added. Samples were incubated on ice for 10 min. After 1 min of shaking, absorbance at 540 nm was taken using Varioskan^TM^ LUX multimode microplate reader. All data was exported into GraphPad Prism^164^ for visualization and analysis. Statistical significance was calculated using unpaired t-test.

## Supporting Item Titles and Legends

**Supplementary Figure 1.**
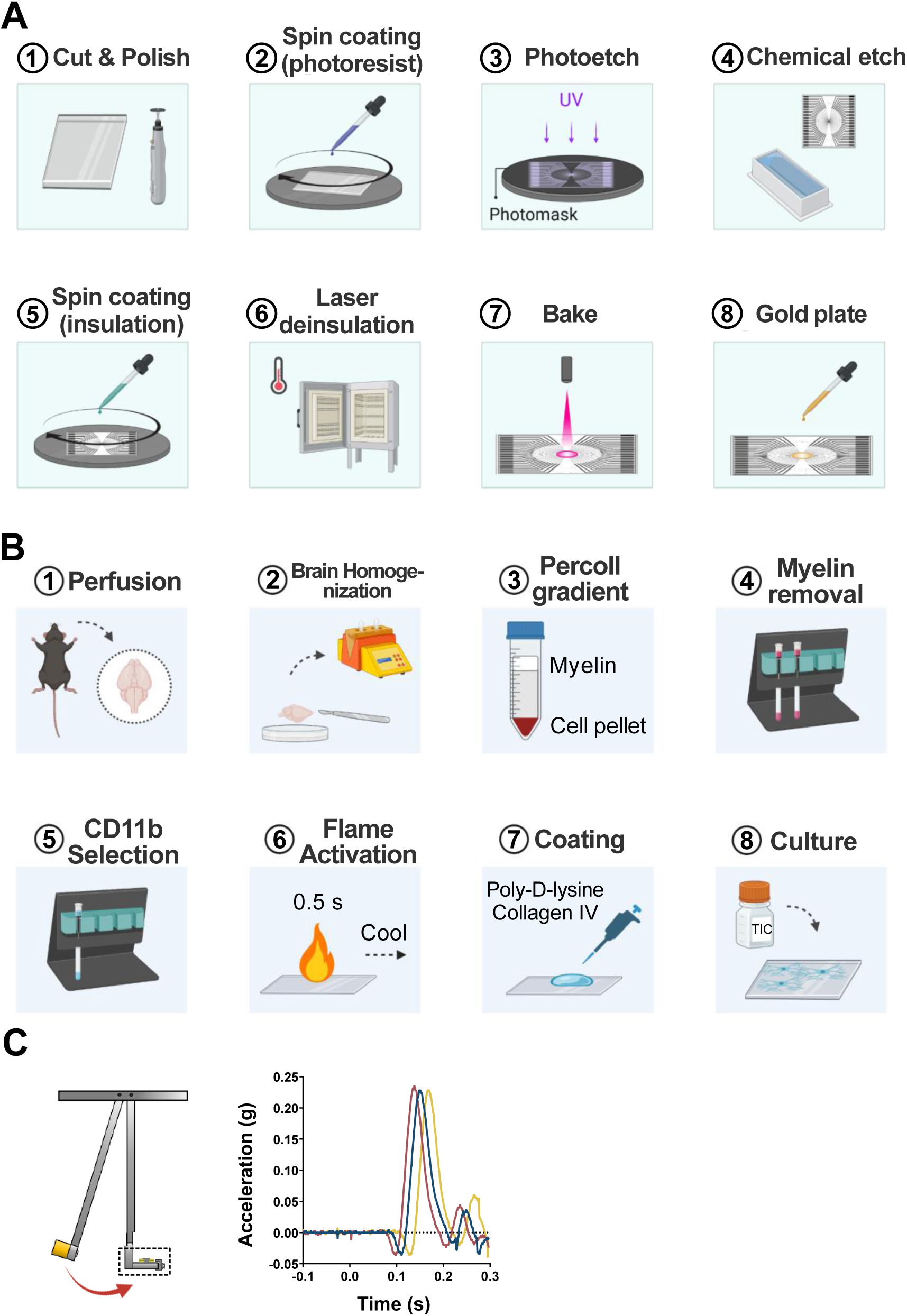
Preparation of glass-based impact chips and primary microglia isolation. **(A)** Scheme of the preparation of multielectrode arrays (MEA, steps 1-8) and enhanced resolution plates (ERP, step 1). **(B)** Scheme of the primary microglia isolation from adult C57 mice. **(C)** Three consecutively sampled and overlaid accelerometer outputs demonstrate the consistency of the impact pendulum.

**Supplementary Figure 2.**
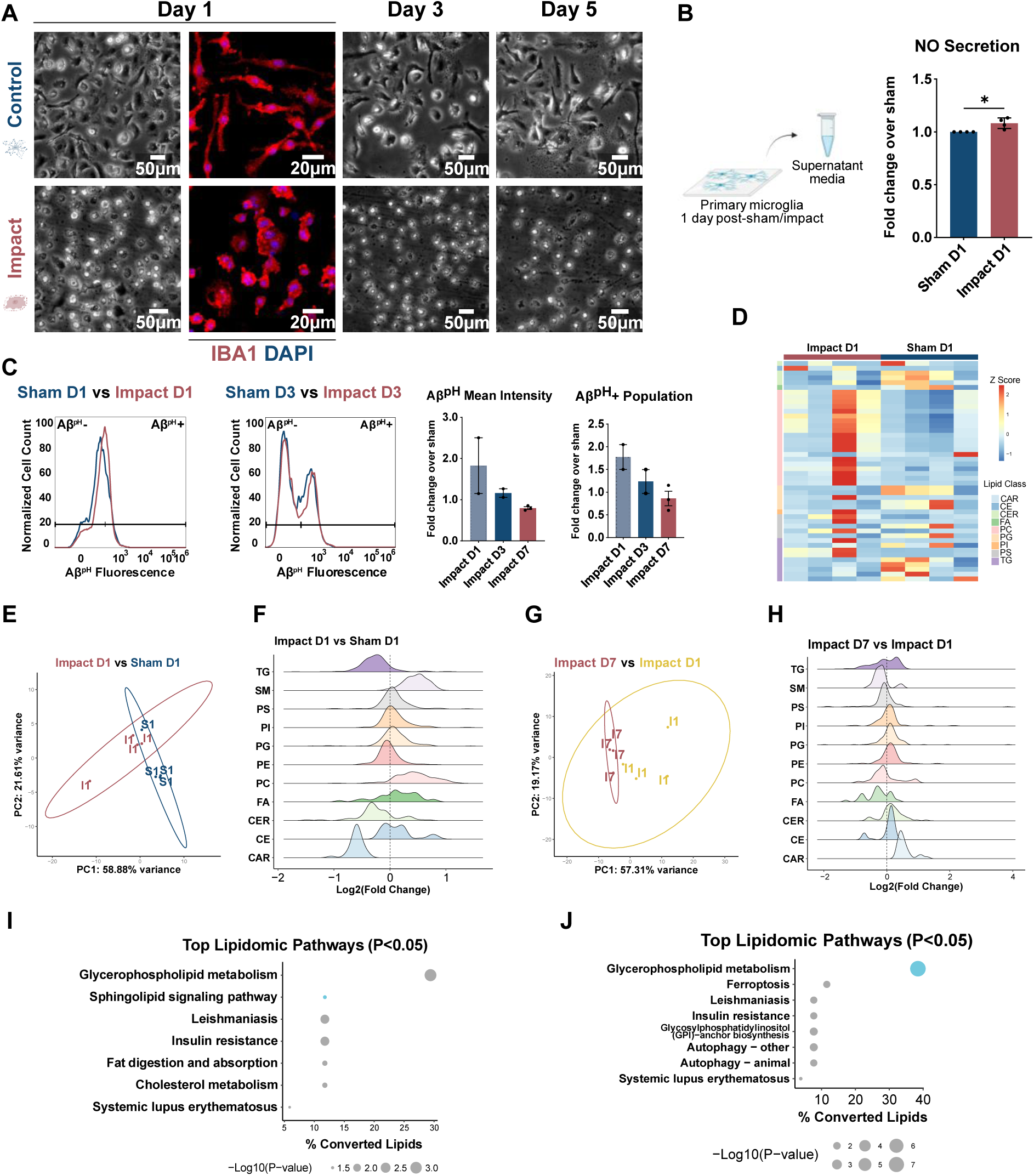
Changes in impacted microglia between 1-7 days post-impact. **(A)** Phase-contrast and fluorescence microscopy images of microglia morphology at 1, 3, and 5 days post-impact (red, IBA1; blue, DAPI). **(B)** Increased NO is significant as early as 1 day post-impact (N=4, p<0.05). **(C)** Decreasing trend in microglial phagocytosis of Aβ^pH^ at 1, 3, and 7 days post-impact measured by flow cytometry (N=2 for Day 1 and 3, N=3 for Day 7). **(D)** Heatmap of significantly changed lipids (FDR < 0.1) at 1 day post-impact. **(E)** PCA and **(F)** Ridge plot of impacted microglial supernatant lipidome at 1 day post-impact vs sham (N=4). **(F)** PCA and **(H)** Ridge plots of impacted microglial supernatant lipidome at 1 day post-impact vs 7 days post-impact. Lipidomic pathway analysis via LIPEA^149^ for significantly changed lipid classes at 1 day post-impact vs **(I) 1 day post-**sham and vs **(J)** 7 days post-impact.

**Supplementary Figure 3.**
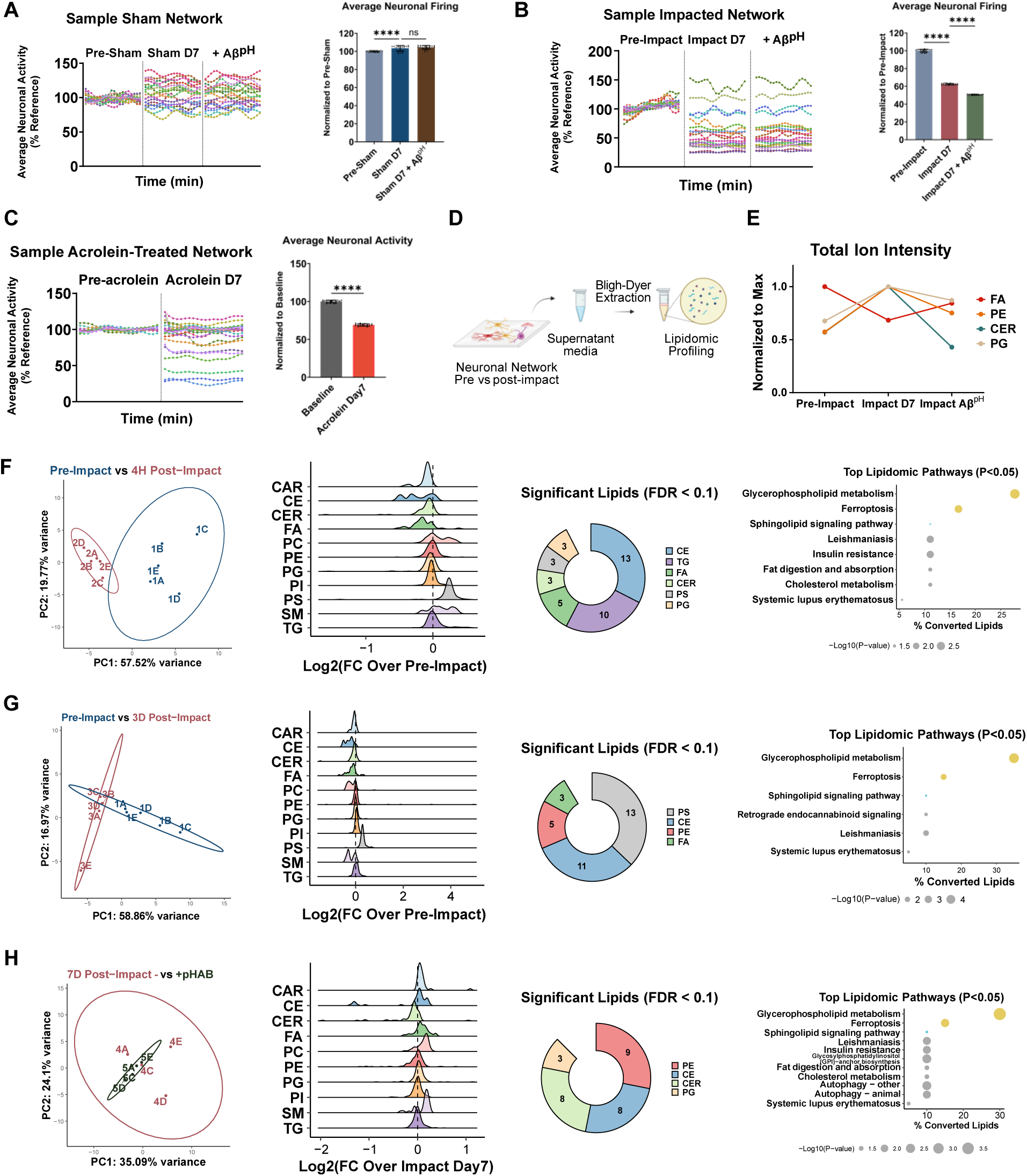
Impact induced electrophysiological and lipidomic changes in neuronal networks at 1-7 days post-impact. Sample electrophysiology results from **(A)** sham, **(B)** impacted, and **(C)** acrolein-treated histiotypical neuronal network. Each dot represents average neuronal activity in one minute, and each line corresponds to one individual neuron measured over 20 minutes. **(D)** Scheme of lipidomic analysis of supernatant media of neuronal networks. PCA analysis of pre-vs 4 hours post-impact (N=5). **(E)** Ridge plot of Log2-tranformed fold change (Impact/Pre-Impact) of all lipids from 11 major lipid classes. **(F)** Composition of significantly changed lipids (FDR < 0.1). **(G)** Total ion intensity of lipid classes whose trends were reversed by Aβ^pH^ treatment (N=4). **(H)** PCA analysis, **(I)** Ridge plot, and **(J)** composition of significant lipids from pre-vs 3 days post-impact (N=5). **(K)** Lipidomic pathway analysis via LIPEA^149^ of 7 days post-impact + vs – Aβ^pH^ treatment. **(L)** PCA analysis, **(M)** Ridge plot, and **(N)** composition of significant lipids from 7 days post-impact + vs – Aβ^pH^ treatment. All error bars indicate SEM. Statistical significance is calculated with Student’s t-test, except for lipidomics data which is calculated with linear regression using the EdgeR package.

**Supplementary Table 1.** Raw lipidomics data of sham and impacted primary microglia supernatant (Sham D1, Sham D7, Impact D1, Impact D7.

**Supplementary Table 2.** Raw lipidomics data of histiotypical neuronal networks supernatant. (Pre-Impact, Impact 4H, Impact D3, Impact D7, Impact D7 + pHAB)

**Supplementary Table 3.** Analyzed lipidomics data of microglia supernatant at 7 days post-sham vs impact.

**Supplementary Table 4.** Analyzed lipidomics data of microglia supernatant at 1 day post-sham vs impact.

**Supplementary Table 5.** Analyzed lipidomics data of microglia supernatant at 1 day vs 7 days post-impact.

**Supplementary Table 6.** Analyzed lipidomics data of neuronal network supernatant at pre-impact vs 4 hours post-impact.

**Supplementary Table 7.** Analyzed lipidomics data of neuronal network supernatant at pre-impact vs 3 days post-impact.

**Supplementary Table 8.** Analyzed lipidomics data of neuronal network supernatant at pre-impact vs 7 days post-impact.

**Supplementary Table 9.** Analyzed lipidomics data of neuronal network supernatant at 7 days post-impact – vs + Aβ^pH^.

**Supplementary Table 10**. BIOPAN pathway analysis of supernatant lipidomic changes in impacted primary microglia.

**Supplementary Table 11.** BIOPAN pathway analysis of supernatant lipidomics changes in impacted neuronal networks.

## Abbreviations

Aβ: amyloid β
Aβ^pH^: pH-responsive Aβ(M1-42)
AD: Alzheimer’s Disease
CAR: acylcarnitine
CE: cholesterol ester, cholesteryl ester
CER: ceramide
CNS: central nervous system
FA: free fatty acid
LPC: lysophosphatidylcholine
NO: nitric oxide
PC: phosphatidylcholine
PE: phosphatidylethanolamine
PG: phosphatidylglycerol
PI: phosphatidylinositol
PS: phosphatidylserine
SM: sphingomyelin
TBI: traumatic brain injury
TG: triacylglycerol

## Notes

https://github.com/chopralab/Impact-Microglia-2024

## References

1. Multiple Cause of Death Data on CDC WONDER https://wonder.cdc.gov/mcd.html.

2. Traumatic brain injury in the United States; emergency department visits, hospitalizations, and deaths, 2002-2006 https://stacks.cdc.gov/view/cdc/5571.

3. Brett, B.L., Gardner, R.C., Godbout, J., Dams-O’Connor, K., and Keene, C.D. (2021). Traumatic Brain Injury and Risk of Neurodegenerative Disorder. Biol. Psychiatry 0. 10.1016/j.biopsych.2021.05.025.

4. Fleminger, S., Oliver, D.L., Lovestone, S., Rabe-Hesketh, S., and Giora, A. (2003). Head injury as a risk factor for Alzheimer’s disease: the evidence 10 years on; a partial replication. J. Neurol. Neurosurg. Psychiatry 74, 857–862. 10.1136/jnnp.74.7.857.

5. Khellaf, A., Khan, D.Z., and Helmy, A. (2019). Recent advances in traumatic brain injury. J. Neurol. 266, 2878–2889. 10.1007/s00415-019-09541-4.

6. Kumaria, A. (2017). In Vitro Models as a Platform to Investigate Traumatic Brain Injury. Altern. Lab. Anim. 45, 201–211. 10.1177/026119291704500405.

7. Menon, D.K. (2009). Unique challenges in clinical trials in traumatic brain injury. Crit. Care Med. 37, S129. 10.1097/CCM.0b013e3181921225.

8. Rogers, E.A., and Gross, G.W. (2019). Simultaneous electrophysiological and morphological assessment of functional damage to neural networks in vitro after 30–300 g impacts. Sci. Rep. 9, 14994. 10.1038/s41598-019-51541-x.

9. Rogers, E.A., Beauclair, T., Thyen, A., and Shi, R. (2022). Utilizing novel TBI-on-a-chip device to link physical impacts to neurodegeneration and decipher primary and secondary injury mechanisms. Sci. Rep. 12, 11838. 10.1038/s41598-022-14937-w.

10. Rogers, E.A., Beauclair, T., Martinez, J., Mufti, S.J., Kim, D., Sun, S., Stingel, R.L., Dieterly, A.M., Krishnan, N., Crodian, J., et al. (2023). The contribution of initial concussive forces and resulting acrolein surge to β-amyloid accumulation and functional alterations in neuronal networks using a TBI-on-a-chip model. Lab. Chip 23, 3388–3404. 10.1039/D3LC00248A.

11. Greve, M.W., and Zink, B.J. (2009). Pathophysiology of traumatic brain injury. Mt. Sinai J. Med. J. Transl. Pers. Med. 76, 97–104. 10.1002/msj.20104.

12. Hamann, K., and Shi, R. (2009). Acrolein scavenging: a potential novel mechanism of attenuating oxidative stress following spinal cord injury. J. Neurochem. 111, 1348–1356. 10.1111/j.1471-4159.2009.06395.x.

13. Kulbe, J.R., and Hall, E.D. (2017). Chronic traumatic encephalopathy-integration of canonical traumatic brain injury secondary injury mechanisms with tau pathology. Prog. Neurobiol. 158, 15–44. 10.1016/j.pneurobio.2017.08.003.

14. Morton, M.V., and Wehman, P. (1995). Psychosocial and emotional sequelae of individuals with traumatic brain injury: a literature review and recommendations. Brain Inj. 9, 81–92. 10.3109/02699059509004574.

15. Barber, C.N., and Raben, D.M. (2019). Lipid Metabolism Crosstalk in the Brain: Glia and Neurons. Front. Cell. Neurosci. 13.

16. Hamilton, J.A., Hillard, C.J., Spector, A.A., and Watkins, P.A. (2007). Brain Uptake and Utilization of Fatty Acids, Lipids and Lipoproteins: Application to Neurological Disorders. J. Mol. Neurosci. 33, 2–11. 10.1007/s12031-007-0060-1.

17. O’Brien, J.S., and Sampson, E.L. (1965). Lipid composition of the normal human brain: gray matter, white matter, and myelin *. J. Lipid Res. 6, 537–544. 10.1016/S0022-2275(20)39619-X.

18. Nessel, I., and Michael-Titus, A.T. (2021). Lipid profiling of brain tissue and blood after traumatic brain injury: A review of human and experimental studies. Semin. Cell Dev. Biol. 112, 145–156. 10.1016/j.semcdb.2020.08.004.

19. Hamann, K., Durkes, A., Ouyang, H., Uchida, K., Pond, A., and Shi, R. (2008). Critical role of acrolein in secondary injury following ex vivo spinal cord trauma. J. Neurochem. 107, 712–721. 10.1111/j.1471-4159.2008.05622.x.

20. Liu-Snyder, P., McNally, H., Shi, R., and Borgens, R.B. (2006). Acrolein-mediated mechanisms of neuronal death. J. Neurosci. Res. 84, 209–218. 10.1002/jnr.20863.

21. Park, J., Muratori, B., and Shi, R. (2014). Acrolein as a novel therapeutic target for motor and sensory deficits in spinal cord injury. Neural Regen. Res. 9, 677–683. 10.4103/1673-5374.131564.

22. Calingasan, N.Y., Uchida, K., and Gibson, G.E. (1999). Protein-Bound Acrolein. J. Neurochem. 72, 751–756. 10.1046/j.1471-4159.1999.0720751.x.

23. Luo, J., Uchida, K., and Shi, R. (2005). Accumulation of Acrolein–Protein Adducts after Traumatic Spinal Cord Injury. Neurochem. Res. 30, 291–295. 10.1007/s11064-005-2602-7.

24. Lawson, L.J., Perry, V.H., Dri, P., and Gordon, S. (1990). Heterogeneity in the distribution and morphology of microglia in the normal adult mouse brain. Neuroscience 39, 151–170. 10.1016/0306-4522(90)90229-W.

25. Shao, F., Wang, X., Wu, H., Wu, Q., and Zhang, J. (2022). Microglia and Neuroinflammation: Crucial Pathological Mechanisms in Traumatic Brain Injury-Induced Neurodegeneration. Front. Aging Neurosci. 14, 825086. 10.3389/fnagi.2022.825086.

26. Davalos, D., Grutzendler, J., Yang, G., Kim, J.V., Zuo, Y., Jung, S., Littman, D.R., Dustin, M.L., and Gan, W.-B. (2005). ATP mediates rapid microglial response to local brain injury in vivo. Nat. Neurosci. 8, 752–758. 10.1038/nn1472.

27. Herzog, C., Pons Garcia, L., Keatinge, M., Greenald, D., Moritz, C., Peri, F., and Herrgen, L. (2019). Rapid clearance of cellular debris by microglia limits secondary neuronal cell death after brain injury in vivo. Development 146, dev174698. 10.1242/dev.174698.

28. Wen, R.-X., Shen, H., Huang, S.-X., Wang, L.-P., Li, Z.-W., Peng, P., Mamtilahun, M., Tang, Y.-H., Shen, F.-X., Tian, H.-L., et al. (2020). P2Y6 receptor inhibition aggravates ischemic brain injury by reducing microglial phagocytosis. CNS Neurosci. Ther. 26, 416–429. 10.1111/cns.13296.

29. Fukumoto, Y., Tanaka, K.F., Parajuli, B., Shibata, K., Yoshioka, H., Kanemaru, K., Gachet, C., Ikenaka, K., Koizumi, S., and Kinouchi, H. (2019). Neuroprotective effects of microglial P2Y1 receptors against ischemic neuronal injury. J. Cereb. Blood Flow Metab. 39, 2144–2156. 10.1177/0271678X18805317.

30. Krukowski, K., Nolan, A., Becker, M., Picard, K., Vernoux, N., Frias, E.S., Feng, X., Tremblay, M.-E., and Rosi, S. (2021). Novel microglia-mediated mechanisms underlying synaptic loss and cognitive impairment after traumatic brain injury. Brain. Behav. Immun. 98, 122–135. 10.1016/j.bbi.2021.08.210.

31. Yip, P.K., Bowes, A.L., Hall, J.C.E., Burguillos, M.A., Ip, T.H.R., Baskerville, T., Liu, Z.-H., Mohamed, M.A.E.K., Getachew, F., Lindsay, A.D., et al. (2019). Docosahexaenoic acid reduces microglia phagocytic activity via miR-124 and induces neuroprotection in rodent models of spinal cord contusion injury. Hum. Mol. Genet. 28, 2427–2448. 10.1093/hmg/ddz073.

32. Butler, C.A., Popescu, A.S., Kitchener, E.J.A., Allendorf, D.H., Puigdellívol, M., and Brown, G.C. (2021). Microglial phagocytosis of neurons in neurodegeneration, and its regulation. J. Neurochem. 158, 621–639. 10.1111/jnc.15327.

33. Chausse, B., Kakimoto, P.A., and Kann, O. (2021). Microglia and lipids: how metabolism controls brain innate immunity. Semin. Cell Dev. Biol. 112, 137–144. 10.1016/j.semcdb.2020.08.001.

34. Prakash, P., Randolph, C.E., Walker, K.A., and Chopra, G. Lipids: Emerging Players of Microglial Biology. Glia n/a. 10.1002/glia.24654.

35. Folick, A., Koliwad, S.K., and Valdearcos, M. (2021). Microglial Lipid Biology in the Hypothalamic Regulation of Metabolic Homeostasis. Front. Endocrinol. 12. 10.3389/fendo.2021.668396.

36. Zhu, X., Zhang, H., Peng, W., Gao, S., Yang, Z., Zhang, J., and Liu, X. (2023). Autophagy impairment is involved in midazolam-induced lipid droplet accumulation and consequent phagocytosis decrease in BV2 cells. Biochem. Biophys. Res. Commun. 643, 147–156. 10.1016/j.bbrc.2022.12.067.

37. Marschallinger, J., Iram, T., Zardeneta, M., Lee, S.E., Lehallier, B., Haney, M.S., Pluvinage, J.V., Mathur, V., Hahn, O., Morgens, D.W., et al. (2020). Lipid-droplet-accumulating microglia represent a dysfunctional and proinflammatory state in the aging brain. Nat. Neurosci. 23, 194–208. 10.1038/s41593-019-0566-1.

38. Nayak, D., Huo, Y., Kwang, W.X.T., Pushparaj, P.N., Kumar, S.D., Ling, E.-A., and Dheen, S.T. (2010). Sphingosine kinase 1 regulates the expression of proinflammatory cytokines and nitric oxide in activated microglia. Neuroscience 166, 132–144. 10.1016/j.neuroscience.2009.12.020.

39. Lu, Z., Liu, S., Lopes-Virella, M.F., and Wang, Z. (2021). LPS and palmitic acid Co-upregulate microglia activation and neuroinflammatory response. Compr. Psychoneuroendocrinology 6, 100048. 10.1016/j.cpnec.2021.100048.

40. Prakash, P., Manchanda, P., Paouri, E., Bisht, K., Sharma, K., Rajpoot, J., Wendt, V., Hossain, A., Wijewardhane, P.R., Randolph, C.E., et al. (2025). Amyloid-β induces lipid droplet-mediated microglial dysfunction via the enzyme DGAT2 in Alzheimer’s disease. Immunity 58, 1536–1552.e8. 10.1016/j.immuni.2025.04.029.

41. Eiserich, J., van der Vliet, A., Handelman, G., Halliwell, B., and Cross, C. (1995). Dietary antioxidants and cigarette smoke-induced biomolecular damage: a complex interaction. Am. J. Clin. Nutr. 62, 1490S–1500S. 10.1093/ajcn/62.6.1490S.

42. Liu-Snyder, P., Logan, M.P., Shi, R., Smith, D.T., and Borgens, R.B. (2007). Neuroprotection from secondary injury by polyethylene glycol requires its internalization. J. Exp. Biol. 210, 1455–1462. 10.1242/jeb.02756.

43. Satoh, K., Yamada, S., Koike, Y., Igarashi, Y., Toyokuni, S., Kumano, T., Takahata, T., Hayakari, M., Tsuchida, S., and Uchida, K. (1999). A 1-Hour Enzyme-Linked Immunosorbent Assay for Quantitation of Acrolein- and Hydroxynonenal-Modified Proteins by Epitope-Bound Casein Matrix Method. Anal. Biochem. 270, 323–328. 10.1006/abio.1999.4073.

44. Fujita, M., Wei, E.P., and Povlishock, J.T. (2012). Intensity- and Interval-Specific Repetitive Traumatic Brain Injury Can Evoke Both Axonal and Microvascular Damage. J. Neurotrauma 29, 2172–2180. 10.1089/neu.2012.2357.

45. Wu, L., Kalish, B.T., Finander, B., Cao, T., Jin, G., Yahya, T., Levy, E.S., Kukreja, B., LaRovere, E.S., Chung, J.Y., et al. (2022). Repetitive Mild Closed Head Injury in Adolescent Mice Is Associated with Impaired Proteostasis, Neuroinflammation, and Tauopathy. J. Neurosci. 42, 2418– 2432. 10.1523/JNEUROSCI.0682-21.2021.

46. Younger, D., Murugan, M., Rama Rao, K.V., Wu, L.-J., and Chandra, N. (2019). Microglia Receptors in Animal Models of Traumatic Brain Injury. Mol. Neurobiol. 56, 5202–5228. 10.1007/s12035-018-1428-7.

47. Orihara, Y., Ikematsu, K., Tsuda, R., and Nakasono, I. (2001). Induction of nitric oxide synthase by traumatic brain injury. Forensic Sci. Int. 123, 142–149. 10.1016/S0379-0738(01)00537-0.

48. Donat, C.K., Scott, G., Gentleman, S.M., and Sastre, M. (2017). Microglial Activation in Traumatic Brain Injury. Front. Aging Neurosci. 9.

49. Garcia-Reitboeck, P., Phillips, A., Piers, T.M., Villegas-Llerena, C., Butler, M., Mallach, A., Rodrigues, C., Arber, C.E., Heslegrave, A., Zetterberg, H., et al. (2018). Human Induced Pluripotent Stem Cell-Derived Microglia-Like Cells Harboring TREM2 Missense Mutations Show Specific Deficits in Phagocytosis. Cell Rep. 24, 2300–2311. 10.1016/j.celrep.2018.07.094.

50. Maguire, E., Connor-Robson, N., Shaw, B., O’Donoghue, R., Stöberl, N., and Hall-Roberts, H. (2022). Assaying Microglia Functions In Vitro. Cells 11, 3414. 10.3390/cells11213414.

51. Prakash, P., P. Jethava, K., Korte, N., Izquierdo, P., Favuzzi, E., L. Rose, I.V., A. Guttenplan, K., Manchanda, P., Dutta, S., Rochet, J.-C., et al. (2021). Monitoring phagocytic uptake of amyloid β into glial cell lysosomes in real time. Chem. Sci. 12, 10901–10918. 10.1039/D1SC03486C.

52. Tejera, D., Mercan, D., Sanchez-Caro, J.M., Hanan, M., Greenberg, D., Soreq, H., Latz, E., Golenbock, D., and Heneka, M.T. (2019). Systemic inflammation impairs microglial Aβ clearance through NLRP3 inflammasome. EMBO J. 38, e101064. 10.15252/embj.2018101064.

53. Bido, S., Muggeo, S., Massimino, L., Marzi, M.J., Giannelli, S.G., Melacini, E., Nannoni, M., Gambarè, D., Bellini, E., Ordazzo, G., et al. (2021). Microglia-specific overexpression of α-synuclein leads to severe dopaminergic neurodegeneration by phagocytic exhaustion and oxidative toxicity. Nat. Commun. 12, 6237. 10.1038/s41467-021-26519-x.

54. Guttenplan, K.A., Weigel, M.K., Prakash, P., Wijewardhane, P.R., Hasel, P., Rufen-Blanchette, U., Münch, A.E., Blum, J.A., Fine, J., Neal, M.C., et al. (2021). Neurotoxic reactive astrocytes induce cell death via saturated lipids. Nature, 1–6. 10.1038/s41586-021-03960-y.

55. Byrns, C.N., Perlegos, A.E., Miller, K.N., Jin, Z., Carranza, F.R., Manchandra, P., Beveridge, C.H., Randolph, C.E., Chaluvadi, V.S., Zhang, S.L., et al. (2024). Senescent glia link mitochondrial dysfunction and lipid accumulation. Nature 630, 475–483. 10.1038/s41586-024-07516-8.

56. Nieweg, K., Schaller, H., and Pfrieger, F.W. (2009). Marked differences in cholesterol synthesis between neurons and glial cells from postnatal rats. J. Neurochem. 109, 125–134. 10.1111/j.1471-4159.2009.05917.x.

57. Ho, W.Y., Hartmann, H., and Ling, S.-C. (2022). Central nervous system cholesterol metabolism in health and disease. IUBMB Life 74, 826–841. 10.1002/iub.2662.

58. Henderson, B.W., Greathouse, K.M., Ramdas, R., Walker, C.K., Rao, T.C., Bach, S.V., Curtis, K.A., Day, J.J., Mattheyses, A.L., and Herskowitz, J.H. (2019). Pharmacologic inhibition of LIMK1 provides dendritic spine resilience against β-amyloid. Sci. Signal. 12, eaaw9318. 10.1126/scisignal.aaw9318.

59. Schulte, S., Gries, M., Christmann, A., and Schäfer, K.-H. (2021). Using multielectrode arrays to investigate neurodegenerative effects of the amyloid-beta peptide. Bioelectron. Med. 7, 15. 10.1186/s42234-021-00078-4.

60. Varghese, K., Molnar, P., Das, M., Bhargava, N., Lambert, S., Kindy, M.S., and Hickman, J.J. (2010). A New Target for Amyloid Beta Toxicity Validated by Standard and High-Throughput Electrophysiology. PLOS ONE 5, e8643. 10.1371/journal.pone.0008643.

61. Jia, Z., Zhu, H., Li, J., Wang, X., Misra, H., and Li, Y. (2012). Oxidative stress in spinal cord injury and antioxidant-based intervention. Spinal Cord 50, 264–274. 10.1038/sc.2011.111.

62. Shi, R., Page, J.C., and Tully, M. (2015). Molecular mechanisms of acrolein-mediated myelin destruction in CNS trauma and disease. Free Radic. Res. 49, 888–895. 10.3109/10715762.2015.1021696.

63. Shi, R., Rickett, T., and Sun, W. (2011). Acrolein-mediated injury in nervous system trauma and diseases. Mol. Nutr. Food Res. 55, 1320–1331. 10.1002/mnfr.201100217.

64. Acosta, G., Race, N., Herr, S., Fernandez, J., Tang, J., Rogers, E., and Shi, R. (2019). Acrolein-mediated alpha-synuclein pathology involvement in the early post-injury pathogenesis of mild blast-induced Parkinsonian neurodegeneration. Mol. Cell. Neurosci. 98, 140–154. 10.1016/j.mcn.2019.06.004.

65. Park, J., Zheng, L., Marquis, A., Walls, M., Duerstock, B., Pond, A., Vega-Alvarez, S., Wang, H., Ouyang, Z., and Shi, R. (2014). Neuroprotective role of hydralazine in rat spinal cord injury-attenuation of acrolein-mediated damage. J. Neurochem. 129, 339–349. 10.1111/jnc.12628.

66. Montgomery, E.A., Fenton, G.W., McClelland, R.J., MacFlynn, G., and Rutherford, W.H. (1991). The psychobiology of minor head injury. Psychol. Med. 21, 375–384. 10.1017/S0033291700020481.

67. Rapp, P.E., Keyser, D.O., Albano, A., Hernandez, R., Gibson, D.B., Zambon, R.A., Hairston, W.D., Hughes, J.D., Krystal, A., and Nichols, A.S. (2015). Traumatic Brain Injury Detection Using Electrophysiological Methods. Front. Hum. Neurosci. 9.

68. Meeting, A. for R. in N. and M.D. (1996). Biology of Schizophrenia and Affective Disease (American Psychiatric Pub).

69. Zhang, Y.P., Cai, J., Shields, L.B.E., Liu, N., Xu, X.-M., and Shields, C.B. (2014). Traumatic Brain Injury Using Mouse Models. Transl. Stroke Res. 5, 454–471. 10.1007/s12975-014-0327-0.

70. Creed, J.A., DiLeonardi, A.M., Fox, D.P., Tessler, A.R., and Raghupathi, R. (2011). Concussive Brain Trauma in the Mouse Results in Acute Cognitive Deficits and Sustained Impairment of Axonal Function. J. Neurotrauma 28, 547–563. 10.1089/neu.2010.1729.

71. Ramlackhansingh, A.F., Brooks, D.J., Greenwood, R.J., Bose, S.K., Turkheimer, F.E., Kinnunen, K.M., Gentleman, S., Heckemann, R.A., Gunanayagam, K., Gelosa, G., et al. (2011). Inflammation after trauma: Microglial activation and traumatic brain injury. Ann. Neurol. 70, 374–383. 10.1002/ana.22455.

72. Yu, F., Wang, Y., Stetler, A.R., Leak, R.K., Hu, X., and Chen, J. (2022). Phagocytic microglia and macrophages in brain injury and repair. CNS Neurosci. Ther. 28, 1279–1293. 10.1111/cns.13899.

73. Fountain, A., Inpanathan, S., Alves, P., Verdawala, M.B., and Botelho, R.J. (2021). Phagosome maturation in macrophages: Eat, digest, adapt, and repeat. Adv. Biol. Regul. 82, 100832. 10.1016/j.jbior.2021.100832.

74. Lee, J.-H., Yang, D.-S., Goulbourne, C.N., Im, E., Stavrides, P., Pensalfini, A., Chan, H., Bouchet-Marquis, C., Bleiwas, C., Berg, M.J., et al. (2022). Faulty autolysosome acidification in Alzheimer’s disease mouse models induces autophagic build-up of Aβ in neurons, yielding senile plaques. Nat. Neurosci. 25, 688–701. 10.1038/s41593-022-01084-8.

75. Lee, J.-H., Yu, W.H., Kumar, A., Lee, S., Mohan, P.S., Peterhoff, C.M., Wolfe, D.M., Martinez-Vicente, M., Massey, A.C., Sovak, G., et al. (2010). Lysosomal Proteolysis and Autophagy Require Presenilin 1 and Are Disrupted by Alzheimer-Related PS1 Mutations. Cell 141, 1146–1158. 10.1016/j.cell.2010.05.008.

76. Gabandé-Rodríguez, E., Keane, L., and Capasso, M. (2020). Microglial phagocytosis in aging and Alzheimer’s disease. J. Neurosci. Res. 98, 284–298. 10.1002/jnr.24419.

77. Alaamery, M., Albesher, N., Aljawini, N., Alsuwailm, M., Massadeh, S., Wheeler, M.A., Chao, C.-C., and Quintana, F.J. (2021). Role of sphingolipid metabolism in neurodegeneration. J. Neurochem. 158, 25–35. 10.1111/jnc.15044.

78. Tamboli, I.Y., Tien, N.T., and Walter, J. (2011). Sphingolipid storage impairs autophagic clearance of Alzheimer-associated proteins. Autophagy 7, 645–646. 10.4161/auto.7.6.15122.

79. Filippov, V., Song, M.A., Zhang, K., Vinters, H.V., Tung, S., Kirsch, W.M., Yang, J., and Duerksen-Hughes, P.J. (2012). Increased Ceramide in Brains with Alzheimer’s and Other Neurodegenerative Diseases. J. Alzheimers Dis. 29, 537–547. 10.3233/JAD-2011-111202.

80. Gier, E.C., Pulliam, A.N., Gaul, D.A., Moore, S.G., LaPlaca, M.C., and Fernández, F.M. (2022). Lipidome Alterations following Mild Traumatic Brain Injury in the Rat. Metabolites 12, 150. 10.3390/metabo12020150.

81. Mondal, K., Takahashi, H., Cole II, J., Mar, N.D., Li, C., Stephenson, D., Chalfant, C., Allegood, J., Cowart, L.A., Reiner, A., et al. (2021). N-3 polyunsaturated fatty acids (n-3 PUFA) prevent traumatic brain injury (TBI)-mediated visual and motor deficits in mice by suppressing ceramide biosynthesis. Invest. Ophthalmol. Vis. Sci. 62, 3030.

82. Lee, S.H., Kho, A.R., Lee, S.H., Hong, D.K., Kang, B.S., Park, M.K., Lee, C.J., Yang, H.W., Woo, S.Y., Park, S.W., et al. (2022). Acid Sphingomyelinase Inhibitor, Imipramine, Reduces Hippocampal Neuronal Death after Traumatic Brain Injury. Int. J. Mol. Sci. 23, 14749. 10.3390/ijms232314749.

83. Chen, C.-S., Rosenwald, A.G., and Pagano, R.E. (1995). Ceramide As a Modulator of Endocytosis ∗. J. Biol. Chem. 270, 13291–13297. 10.1074/jbc.270.22.13291.

84. Hjorth, E., Zhu, M., Toro, V.C., Vedin, I., Palmblad, J., Cederholm, T., Freund-Levi, Y., Faxen-Irving, G., Wahlund, L.-O., Basun, H., et al. (2013). Omega-3 Fatty Acids Enhance Phagocytosis of Alzheimer’s Disease-Related Amyloid-β 42 by Human Microglia and Decrease Inflammatory Markers. J. Alzheimers Dis. 35, 697–713. 10.3233/JAD-130131.

85. Chen, S., Zhang, H., Pu, H., Wang, G., Li, W., Leak, R.K., Chen, J., Liou, A.K., and Hu, X. (2014). n-3 PUFA supplementation benefits microglial responses to myelin pathology. Sci. Rep. 4, 7458. 10.1038/srep07458.

86. Tracy, L.M., Bergqvist, F., Ivanova, E.V., Jacobsen, K.T., and Iverfeldt, K. (2013). Exposure to the Saturated Free Fatty Acid Palmitate Alters BV-2 Microglia Inflammatory Response. J. Mol. Neurosci. 51, 805–812. 10.1007/s12031-013-0068-7.

87. Nadjar, A. (2018). Role of metabolic programming in the modulation of microglia phagocytosis by lipids. Prostaglandins Leukot. Essent. Fatty Acids 135, 63–73. 10.1016/j.plefa.2018.07.006.

88. Yanguas-Casás, N., Crespo-Castrillo, A., de Ceballos, M.L., Chowen, J.A., Azcoitia, I., Arevalo, M.A., and Garcia-Segura, L.M. (2018). Sex differences in the phagocytic and migratory activity of microglia and their impairment by palmitic acid. Glia 66, 522–537. 10.1002/glia.23263.

89. Chandak, P.G., Radović, B., Aflaki, E., Kolb, D., Buchebner, M., Fröhlich, E., Magnes, C., Sinner, F., Haemmerle, G., Zechner, R., et al. (2010). Efficient Phagocytosis Requires Triacylglycerol Hydrolysis by Adipose Triglyceride Lipase*. J. Biol. Chem. 285, 20192–20201. 10.1074/jbc.M110.107854.

90. Huang, S.C.-C., Everts, B., Ivanova, Y., O’Sullivan, D., Nascimento, M., Smith, A.M., Beatty, W., Love-Gregory, L., Lam, W.Y., O’Neill, C.M., et al. (2014). Cell-intrinsic lysosomal lipolysis is essential for alternative activation of macrophages. Nat. Immunol. 15, 846–855. 10.1038/ni.2956.

91. Birge, R.B., Boeltz, S., Kumar, S., Carlson, J., Wanderley, J., Calianese, D., Barcinski, M., Brekken, R.A., Huang, X., Hutchins, J.T., et al. (2016). Phosphatidylserine is a global immunosuppressive signal in efferocytosis, infectious disease, and cancer. Cell Death Differ. 23, 962–978. 10.1038/cdd.2016.11.

92. Freikman, I., Ringel, I., and Fibach, E. (2011). Oxidative Stress-Induced Membrane Shedding from RBCs is Ca Flux-Mediated and Affects Membrane Lipid Composition. J. Membr. Biol. 240, 73–82. 10.1007/s00232-011-9345-y.

93. Ma, X., Li, X., Wang, W., Zhang, M., Yang, B., and Miao, Z. (2022). Phosphatidylserine, inflammation, and central nervous system diseases. Front. Aging Neurosci. 14.

94. Huang, L., Tang, H., Sherchan, P., Lenahan, C., Boling, W., Tang, J., and Zhang, J.H. (2020). The Activation of Phosphatidylserine/CD36/TGF-β1 Pathway prior to Surgical Brain Injury Attenuates Neuroinflammation in Rats. Oxid. Med. Cell. Longev. 2020, 4921562. 10.1155/2020/4921562.

95. Mallah, K., Quanico, J., Raffo-Romero, A., Cardon, T., Aboulouard, S., Devos, D., Kobeissy, F., Zibara, K., Salzet, M., and Fournier, I. (2019). Matrix-Assisted Laser Desorption/Ionization-Mass Spectrometry Imaging of Lipids in Experimental Model of Traumatic Brain Injury Detecting Acylcarnitines as Injury Related Markers. Anal. Chem. 91, 11879–11887. 10.1021/acs.analchem.9b02633.

96. Scafidi, S., Racz, J., Hazelton, J., McKenna, M.C., and Fiskum, G. (2011). Neuroprotection by Acetyl-L-Carnitine after Traumatic Injury to the Immature Rat Brain. Dev. Neurosci. 32, 480–487. 10.1159/000323178.

97. Zhang, R., Zhang, H., Zhang, Z., Wang, T., Niu, J., Cui, D., and Xu, S. (2012). Neuroprotective Effects of Pre-Treament with l-Carnitine and Acetyl-l-Carnitine on Ischemic Injury In Vivo and In Vitro. Int. J. Mol. Sci. 13, 2078–2090. 10.3390/ijms13022078.

98. Jones, L.L., McDonald, D.A., and Borum, P.R. (2010). Acylcarnitines: Role in brain. Prog. Lipid Res. 49, 61–75. 10.1016/j.plipres.2009.08.004.

99. Pettegrew, J.W., Levine, J., and McClure, R.J. (2000). Acetyl-L-carnitine physical-chemical, metabolic, and therapeutic properties: relevance for its mode of action in Alzheimer’s disease and geriatric depression. Mol. Psychiatry 5, 616–632. 10.1038/sj.mp.4000805.

100. Montgomery, S.A., Thal, L.J., and Amrein, R. (2003). Meta-analysis of double blind randomized controlled clinical trials of acetyl-L-carnitine versus placebo in the treatment of mild cognitive impairment and mild Alzheimer’s disease. Int. Clin. Psychopharmacol. 18, 61–71. 10.1097/00004850-200303000-00001.

101. Jones, L.L., McDonald, D.A., and Borum, P.R. (2010). Acylcarnitines: Role in brain. Prog. Lipid Res. 49, 61–75. 10.1016/j.plipres.2009.08.004.

102. Tzekov, R., Dawson, C., Orlando, M., Mouzon, B., Reed, J., Evans, J., Crynen, G., Mullan, M., and Crawford, F. (2016). Sub-Chronic Neuropathological and Biochemical Changes in Mouse Visual System after Repetitive Mild Traumatic Brain Injury. PLoS ONE 11, e0153608. 10.1371/journal.pone.0153608.

103. Koizumi, S., Yamamoto, S., Hayasaka, T., Konishi, Y., Yamaguchi-Okada, M., Goto-Inoue, N., Sugiura, Y., Setou, M., and Namba, H. (2010). Imaging mass spectrometry revealed the production of lyso-phosphatidylcholine in the injured ischemic rat brain. Neuroscience 168, 219–225. 10.1016/j.neuroscience.2010.03.056.

104. Sheikh, A. Md., Michikawa, M., Kim, S.U., and Nagai, A. (2015). Lysophosphatidylcholine increases the neurotoxicity of Alzheimer’s amyloid β1-42 peptide: Role of oligomer formation. Neuroscience 292, 159–169. 10.1016/j.neuroscience.2015.02.034.

105. Kaya, I., Brinet, D., Michno, W., Başkurt, M., Zetterberg, H., Blenow, K., and Hanrieder, J. (2017). Novel Trimodal MALDI Imaging Mass Spectrometry (IMS3) at 10 μm Reveals Spatial Lipid and Peptide Correlates Implicated in Aβ Plaque Pathology in Alzheimer’s Disease. ACS Chem. Neurosci. 8, 2778–2790. 10.1021/acschemneuro.7b00314.

106. Dai, L., Zou, L., Meng, L., Qiang, G., Yan, M., and Zhang, Z. (2021). Cholesterol Metabolism in Neurodegenerative Diseases: Molecular Mechanisms and Therapeutic Targets. Mol. Neurobiol. 58, 2183–2201. 10.1007/s12035-020-02232-6.

107. Essayan-Perez, S., and Südhof, T.C. (2023). Neuronal γ-secretase regulates lipid metabolism, linking cholesterol to synaptic dysfunction in Alzheimer’s disease. Neuron. 10.1016/j.neuron.2023.07.005.

108. Canolle, B., Masmejean, F., Melon, C., Nieoullon, A., Pisano, P., and Lortet, S. (2004). Glial soluble factors regulate the activity and expression of the neuronal glutamate transporter EAAC1: implication of cholesterol. J. Neurochem. 88, 1521–1532. 10.1046/j.1471-4159.2003.02301.x.

109. Sponne, I., Fifre, A., Kriem, B., Koziel, V., Bihain, B., Oster, T., Olivier, J.-L., and Pillot, T. (2004). Membrane cholesterol interferes with neuronal apoptosis induced by soluble oligomers but not fibrils of the amyloid-β peptide. FASEB J. 18, 836–838. 10.1096/fj.03-0372fje.

110. Petrov, A.M., Kasimov, M.R., and Zefirov, A.L. (2016). Brain Cholesterol Metabolism and Its Defects: Linkage to Neurodegenerative Diseases and Synaptic Dysfunction. Acta Naturae 8, 58–73.

111. Luo, J., Yang, H., and Song, B.-L. (2020). Mechanisms and regulation of cholesterol homeostasis. Nat. Rev. Mol. Cell Biol. 21, 225–245. 10.1038/s41580-019-0190-7.

112. Wang, Y.-J., Bian, Y., Luo, J., Lu, M., Xiong, Y., Guo, S.-Y., Yin, H.-Y., Lin, X., Li, Q., Chang, C.C.Y., et al. (2017). Cholesterol and fatty acids regulate cysteine ubiquitylation of ACAT2 through competitive oxidation. Nat. Cell Biol. 19, 808–819. 10.1038/ncb3551.

113. Kim, J.-H., Ee, S.-M., Jittiwat, J., Ong, E.-S., Farooqui, A.A., Jenner, A.M., and Ong, W.-Y. (2011). Increased expression of acyl-coenzyme A: cholesterol acyltransferase-1 and elevated cholesteryl esters in the hippocampus after excitotoxic injury. Neuroscience 185, 125–134. 10.1016/j.neuroscience.2011.04.018.

114. Chang, T.Y., Chang, C.C.Y., Harned, T.C., De La Torre, A.L., Lee, J., Huynh, T.N., and Gow, J.G. (2021). Blocking cholesterol storage to treat Alzheimer’s disease. Explor. Neuroprotective Ther. 1, 173–184. 10.37349/ent.2021.00014.

115. Phillips, G.R., Hancock, S.E., Brown, S.H.J., Jenner, A.M., Kreilaus, F., Newell, K.A., and Mitchell, T.W. (2020). Cholesteryl ester levels are elevated in the caudate and putamen of Huntington’s disease patients. Sci. Rep. 10, 20314. 10.1038/s41598-020-76973-8.

116. Jha, M.K., and Morrison, B.M. (2018). Glia-neuron energy metabolism in health and diseases: New insights into the role of nervous system metabolic transporters. Exp. Neurol. 309, 23–31. 10.1016/j.expneurol.2018.07.009.

117. Xu, X.-J., Yang, M.-S., Zhang, B., Niu, F., Dong, J.-Q., and Liu, B.-Y. (2021). Glucose metabolism: A link between traumatic brain injury and Alzheimer’s disease. Chin. J. Traumatol. 24, 5–10. 10.1016/j.cjtee.2020.10.001.

118. Cunnane, S.C., Trushina, E., Morland, C., Prigione, A., Casadesus, G., Andrews, Z.B., Beal, M.F., Bergersen, L.H., Brinton, R.D., de la Monte, S., et al. (2020). Brain energy rescue: an emerging therapeutic concept for neurodegenerative disorders of ageing. Nat. Rev. Drug Discov. 19, 609–633. 10.1038/s41573-020-0072-x.

119. Balabanov, R., Goldman, H., Murphy, S., Pellizon, G., Owen, C., Rafols, J., and Dore-Duffy, P. (2001). Endothelial cell activation following moderate traumatic brain injury. Neurol. Res. 23, 175–182. 10.1179/016164101101198514.

120. Cornford, E.M., Hyman, S., Cornford, M.E., and Caron, M.J. (1996). Glut1 glucose transporter activity in human brain injury. J. Neurotrauma 13, 523–536. 10.1089/neu.1996.13.523.

121. Kyrtata, N., Emsley, H.C.A., Sparasci, O., Parkes, L.M., and Dickie, B.R. (2021). A Systematic Review of Glucose Transport Alterations in Alzheimer’s Disease. Front. Neurosci. 15.

122. Lauro, C., and Limatola, C. (2020). Metabolic Reprograming of Microglia in the Regulation of the Innate Inflammatory Response. Front. Immunol. 11.

123. Ashrafi, G., Wu, Z., Farrell, R.J., and Ryan, T.A. (2017). GLUT4 Mobilization Supports Energetic Demands of Active Synapses. Neuron 93, 606–615.e3. 10.1016/j.neuron.2016.12.020.

124. Summers, S.A., Garza, L.A., Zhou, H., and Birnbaum, M.J. (1998). Regulation of Insulin-Stimulated Glucose Transporter GLUT4 Translocation and Akt Kinase Activity by Ceramide. Mol. Cell. Biol. 18, 5457–5464. 10.1128/MCB.18.9.5457.

125. Randle, P.J. (1998). Regulatory interactions between lipids and carbohydrates: the glucose fatty acid cycle after 35 years. Diabetes. Metab. Rev. 14, 263–283. 10.1002/(SICI)1099-0895(199812)14:4.0.CO;2-C.

126. Kagan, V.E., Mao, G., Qu, F., Angeli, J.P.F., Doll, S., Croix, C.S., Dar, H.H., Liu, B., Tyurin, V.A., Ritov, V.B., et al. (2017). Oxidized Arachidonic/Adrenic Phosphatidylethanolamines Navigate Cells to Ferroptosis. Nat. Chem. Biol. 13, 81–90. 10.1038/nchembio.2238.

127. Kenny, E.M., Fidan, E., Yang, Q., Anthonymuthu, T.S., New, L.A., Meyer, E.A., Wang, H., Kochanek, P.M., Dixon, C.E., Kagan, V.E., et al. (2019). Ferroptosis contributes to neuronal death and functional outcome after traumatic brain injury. Crit. Care Med. 47, 410–418. 10.1097/CCM.0000000000003555.

128. Uderhardt, S., Herrmann, M., Oskolkova, O.V., Aschermann, S., Bicker, W., Ipseiz, N., Sarter, K., Frey, B., Rothe, T., Voll, R., et al. (2012). 12/15-Lipoxygenase Orchestrates the Clearance of Apoptotic Cells and Maintains Immunologic Tolerance. Immunity 36, 834–846. 10.1016/j.immuni.2012.03.010.

129. Ansari, M.A., Roberts, K.N., and Scheff, S.W. (2008). Oxidative stress and modification of synaptic proteins in hippocampus after traumatic brain injury. Free Radic. Biol. Med. 45, 443–452. 10.1016/j.freeradbiomed.2008.04.038.

130. Ho, W.-C., Hsu, C.-C., Huang, H.-J., Wang, H.-T., and Lin, A.M.-Y. (2020). Anti-inflammatory Effect of AZD6244 on Acrolein-Induced Neuroinflammation. Mol. Neurobiol. 57, 88–95. 10.1007/s12035-019-01758-8.

131. Huang, H.-J., Wang, H.-T., Yeh, T.-Y., Lin, B.-W., Shiao, Y.-J., Lo, Y.-L., and Lin, A.M.-Y. (2021). Neuroprotective effect of selumetinib on acrolein-induced neurotoxicity. Sci. Rep. 11, 12497. 10.1038/s41598-021-91507-6.

132. Spaas, J., Franssen, W.M.A., Keytsman, C., Blancquaert, L., Vanmierlo, T., Bogie, J., Broux, B., Hellings, N., van Horssen, J., Posa, D.K., et al. (2021). Carnosine quenches the reactive carbonyl acrolein in the central nervous system and attenuates autoimmune neuroinflammation. J. Neuroinflammation 18, 255. 10.1186/s12974-021-02306-9.

133. Khoramjouy, M., Naderi, N., Kobarfard, F., Heidarli, E., and Faizi, M. (2021). An Intensified Acrolein Exposure Can Affect Memory and Cognition in Rat. Neurotox. Res. 39, 277–291. 10.1007/s12640-020-00278-x.

134. Shi, Y., Sun, W., McBride, J.J., Cheng, J.-X., and Shi, R. (2011). Acrolein induces myelin damage in mammalian spinal cord. J. Neurochem. 117, 554–564. 10.1111/j.1471-4159.2011.07226.x.

135. Yan, R., Page, J.C., and Shi, R. (2016). Acrolein-mediated conduction loss is partially restored by K+ channel blockers. J. Neurophysiol. 115, 701–710. 10.1152/jn.00467.2015.

136. Krohne, T.U., Kaemmerer, E., Holz, F.G., and Kopitz, J. (2010). Lipid peroxidation products reduce lysosomal protease activities in human retinal pigment epithelial cells via two different mechanisms of action. Exp. Eye Res. 90, 261–266. 10.1016/j.exer.2009.10.014.

137. Aguilar-Gaytan, R., and Mas-Oliva, J. (2003). Oxidative stress impairs endocytosis of the scavenger receptor class A. Biochem. Biophys. Res. Commun. 305, 510–517. 10.1016/S0006-291X(03)00796-4.

138. Ohnishi, Y., Tsuji, D., and Itoh, K. (2022). Oxidative Stress Impairs Autophagy *via* Inhibition of Lysosomal Transport of VAMP8. Biol. Pharm. Bull. 45, 1609–1615. 10.1248/bpb.b22-00131.

139. Ahmed, M.S.E., Langer, H., Abed, M., Voelkl, J., and Lang, F. (2013). The Uremic Toxin Acrolein Promotes Suicidal Erythrocyte Death. Kidney Blood Press. Res. 37, 158–167. 10.1159/000350141.

140. Wiernicki, B., Dubois, H., Tyurina, Y.Y., Hassannia, B., Bayir, H., Kagan, V.E., Vandenabeele, P., Wullaert, A., and Vanden Berghe, T. (2020). Excessive phospholipid peroxidation distinguishes ferroptosis from other cell death modes including pyroptosis. Cell Death Dis. 11, 1–11. 10.1038/s41419-020-03118-0.

141. Yang, W.S., and Stockwell, B.R. (2016). Ferroptosis: Death by Lipid Peroxidation. Trends Cell Biol. 26, 165–176. 10.1016/j.tcb.2015.10.014.

142. Xie, B.-S., Wang, Y.-Q., Lin, Y., Mao, Q., Feng, J.-F., Gao, G.-Y., and Jiang, J.-Y. (2019). Inhibition of ferroptosis attenuates tissue damage and improves long-term outcomes after traumatic brain injury in mice. CNS Neurosci. Ther. 25, 465–475. 10.1111/cns.13069.

143. Ryan, S.K., Zelic, M., Han, Y., Teeple, E., Chen, L., Sadeghi, M., Shankara, S., Guo, L., Li, C., Pontarelli, F., et al. (2023). Microglia ferroptosis is regulated by SEC24B and contributes to neurodegeneration. Nat. Neurosci. 26, 12–26. 10.1038/s41593-022-01221-3.

144. Kapralov, A.A., Yang, Q., Dar, H.H., Tyurina, Y.Y., Anthonymuthu, T.S., Kim, R., St. Croix, C.M., Mikulska-Ruminska, K., Liu, B., Shrivastava, I.H., et al. (2020). Redox lipid reprogramming commands susceptibility of macrophages and microglia to ferroptotic death. Nat. Chem. Biol. 16, 278–290. 10.1038/s41589-019-0462-8.

145. Singh, R., Kaushik, S., Wang, Y., Xiang, Y., Novak, I., Komatsu, M., Tanaka, K., Cuervo, A.M., and Czaja, M.J. (2009). Autophagy regulates lipid metabolism. Nature 458, 1131–1135. 10.1038/nature07976.

146. Pierzynowska, K., Rintz, E., Gaffke, L., and Węgrzyn, G. (2021). Ferroptosis and Its Modulation by Autophagy in Light of the Pathogenesis of Lysosomal Storage Diseases. Cells 10, 365. 10.3390/cells10020365.

147. Hegdekar, N., Sarkar, C., Bustos, S., Ritzel, R.M., Hanscom, M., Ravishankar, P., Philkana, D., Wu, J., Loane, D.J., and Lipinski, M.M. (2023). Inhibition of autophagy in microglia and macrophages exacerbates innate immune responses and worsens brain injury outcomes. Autophagy 19, 2026– 2044. 10.1080/15548627.2023.2167689.

148. Choi, I., Wang, M., Yoo, S., Xu, P., Seegobin, S.P., Li, X., Han, X., Wang, Q., Peng, J., Zhang, B., et al. (2023). Autophagy enables microglia to engage amyloid plaques and prevents microglial senescence. Nat. Cell Biol. 25, 963–974. 10.1038/s41556-023-01158-0.

149. LIPEA: Lipid Pathway Enrichment Analysis | bioRxiv https://www.biorxiv.org/content/10.1101/274969v1.abstract.

150. Acevedo, A., Durán, C., Ciucci, S., Gerl, M., and Cannistraci, C.V. (2018). LIPEA: Lipid Pathway Enrichment Analysis. Preprint at bioRxiv, 10.1101/274969 https://doi.org/10.1101/274969.

151. Robinson, M.D., McCarthy, D.J., and Smyth, G.K. (2010). edgeR: a Bioconductor package for differential expression analysis of digital gene expression data. Bioinforma. Oxf. Engl. 26, 139–140. 10.1093/bioinformatics/btp616.

152. Scheiblich, H., Schlütter, A., Golenbock, D.T., Latz, E., Martinez-Martinez, P., and Heneka, M.T. (2017). Activation of the NLRP3 inflammasome in microglia: the role of ceramide. J. Neurochem. 143, 534–550. 10.1111/jnc.14225.

153. Corrêa, R., Silva, L.F.F., Ribeiro, D.J.S., Almeida, R. das N., Santos, I. de O., Corrêa, L.H., de Sant’Ana, L.P., Assunção, L.S., Bozza, P.T., and Magalhães, K.G. (2020). Lysophosphatidylcholine Induces NLRP3 Inflammasome-Mediated Foam Cell Formation and Pyroptosis in Human Monocytes and Endothelial Cells. Front. Immunol. 10.

154. Martín, M.G., Pfrieger, F., and Dotti, C.G. (2014). Cholesterol in brain disease: sometimes determinant and frequently implicated. EMBO Rep. 15, 1036–1052. 10.15252/embr.201439225.

155. Lucas, J.H., Czisny, L.E., and Gross, G.W. (1986). Adhesion of cultured mammalian central nervous system neurons to flame-modified hydrophobic surfaces. In Vitro Cell. Dev. Biol. 22, 37–43. 10.1007/BF02623439.

156. Gross, G.W., and Schwalm, F.U. (1994). A closed flow chamber for long-term multichannel recording and optical monitoring. J. Neurosci. Methods 52, 73–85. 10.1016/0165-0270(94)90059-0.

157. Keefer, E.W., Gramowski, A., and Gross, G.W. (2001). NMDA receptor-dependent periodic oscillations in cultured spinal cord networks. J. Neurophysiol. 86, 3030–3042. 10.1152/jn.2001.86.6.3030.

158. Ransom, B.R., Neale, E., Henkart, M., Bullock, P.N., and Nelson, P.G. (1977). Mouse spinal cord in cell culture. I. Morphology and intrinsic neuronal electrophysiologic properties. J. Neurophysiol. 40, 1132–1150. 10.1152/jn.1977.40.5.1132.

159. Gross, G.W., and Gopal, K.V. (2006). Emerging Histiotypic Properties of Cultured Neuronal Networks. In Advances in Network Electrophysiology: Using Multi-Electrode Arrays, M. Taketani and M. Baudry, eds. (Springer US), pp. 193–214. 10.1007/0-387-25858-2_8.

160. Collins, H.Y., and Bohlen, C.J. (2018). Isolation and Culture of Rodent Microglia to Promote a Dynamic Ramified Morphology in Serum-free Medium. JoVE J. Vis. Exp., e57122. 10.3791/57122.

161. Gross, G. (1995). The use of neuronal networks on multielectrode arrays as biosensors. Biosens. Bioelectron. 10, 553.

162. Morefield, S.I., Keefer, E.W., Chapman, K.D., and Gross, G.W. (2000). Drug evaluations using neuronal networks cultured on microelectrode arrays. Biosens. Bioelectron. 15, 383–396. 10.1016/s0956-5663(00)00095-6.

163. Novo, M., Freire, S., and Al-Soufi, W. (2018). Critical aggregation concentration for the formation of early Amyloid-β (1-42) oligomers. Sci. Rep. 8, 1783. 10.1038/s41598-018-19961-3.

164. GraphPad Prism Version 8.0.0.

165. Prakash, P., Lantz, T.C., Jethava, K.P., and Chopra, G. (2019). Rapid, Refined, and Robust Method for Expression, Purification, and Characterization of Recombinant Human Amyloid beta 1-42. Methods Protoc. 2, 48. 10.3390/mps2020048.

166. Schindelin, J., Arganda-Carreras, I., Frise, E., Kaynig, V., Longair, M., Pietzsch, T., Preibisch, S., Rueden, C., Saalfeld, S., Schmid, B., et al. (2012). Fiji: an open-source platform for biological-image analysis. Nat. Methods 9, 676–682. 10.1038/nmeth.2019.

167. Stirling, D.R., Swain-Bowden, M.J., Lucas, A.M., Carpenter, A.E., Cimini, B.A., and Goodman, A. (2021). CellProfiler 4: improvements in speed, utility and usability. BMC Bioinformatics 22, 433. 10.1186/s12859-021-04344-9.

168. FlowJo^TM^ Software (2023). Version 10 (Becton, Dickinson and Company).

169. Ghilarducci, D.P., and Tjeerdema, R.S. (1995). Fate and effects of acrolein. Rev. Environ. Contam. Toxicol. 144, 95–146. 10.1007/978-1-4612-2550-8_2.

170. McCarthy, D.J., Chen, Y., and Smyth, G.K. (2012). Differential expression analysis of multifactor RNA-Seq experiments with respect to biological variation. Nucleic Acids Res. 40, 4288–4297. 10.1093/nar/gks042.

171. Benjamini, Y., and Hochberg, Y. (1995). Controlling the False Discovery Rate: A Practical and Powerful Approach to Multiple Testing. J. R. Stat. Soc. Ser. B Methodol. 57, 289–300. 10.1111/j.2517-6161.1995.tb02031.x.

