## Supplementary figures and images for "Impact Induces Phagocytic Defect in Reactive Microglia"

### Supplemental Figure 1

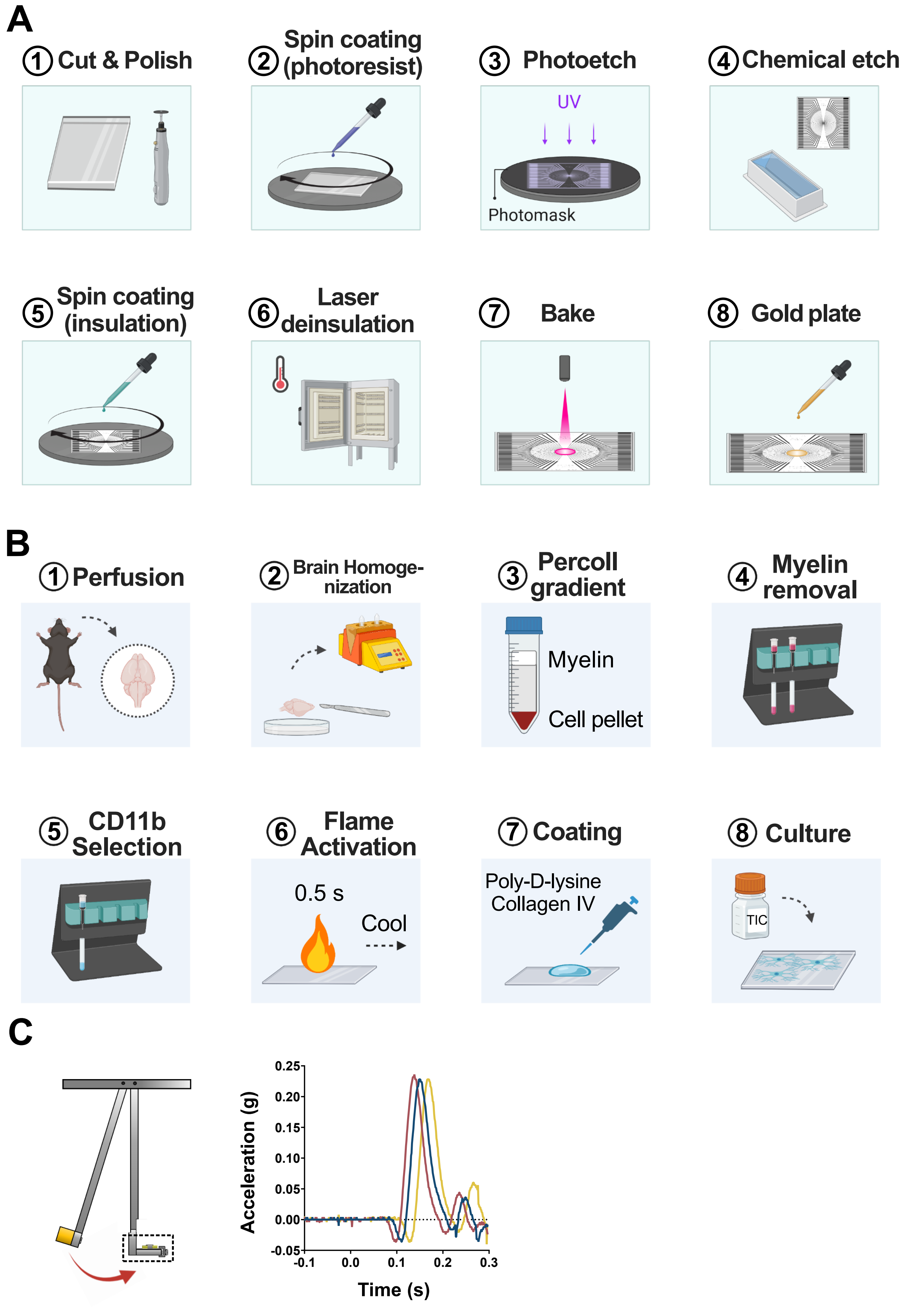

### Supplemental Figure 2

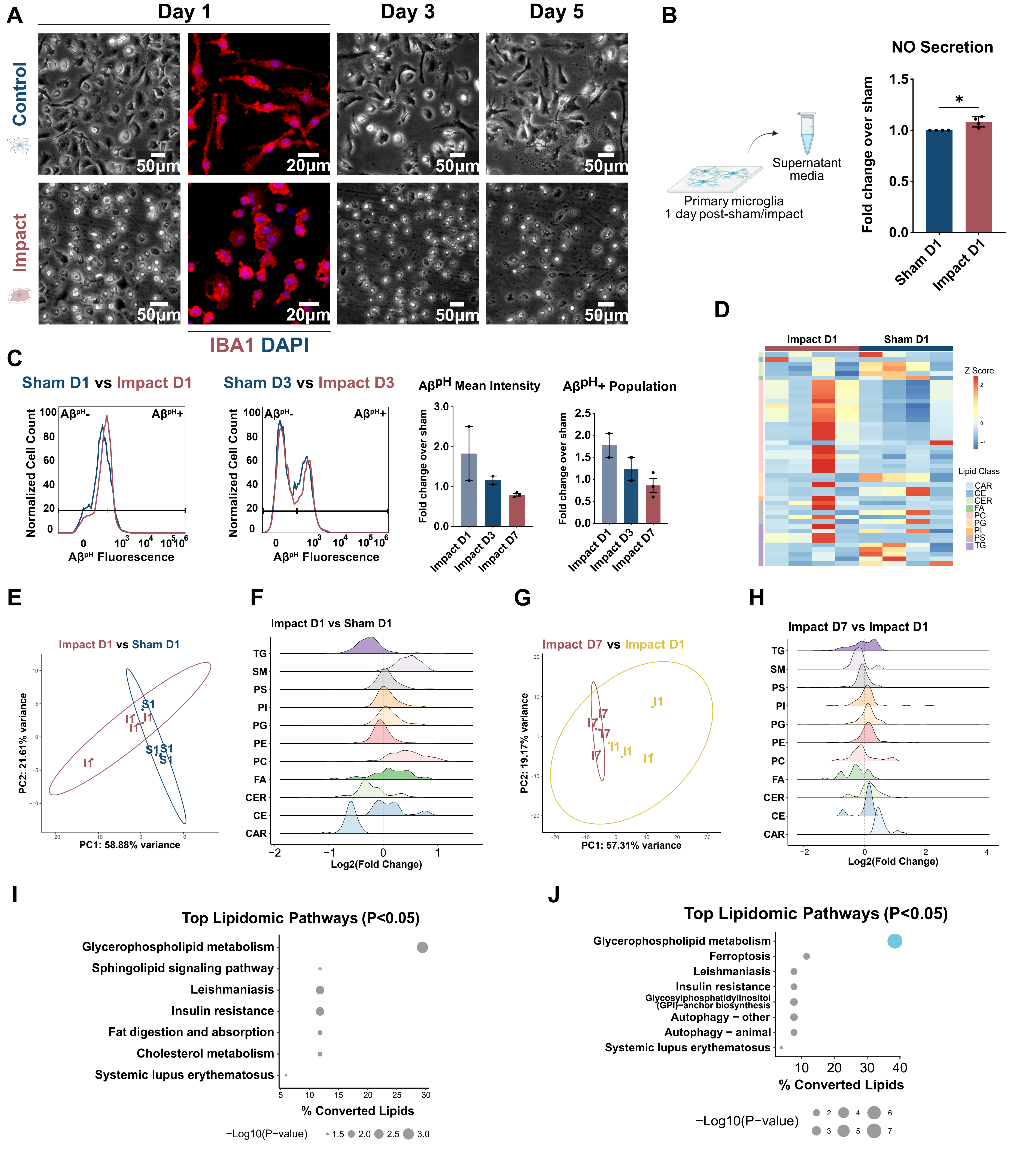

### Supplemental Figure 3

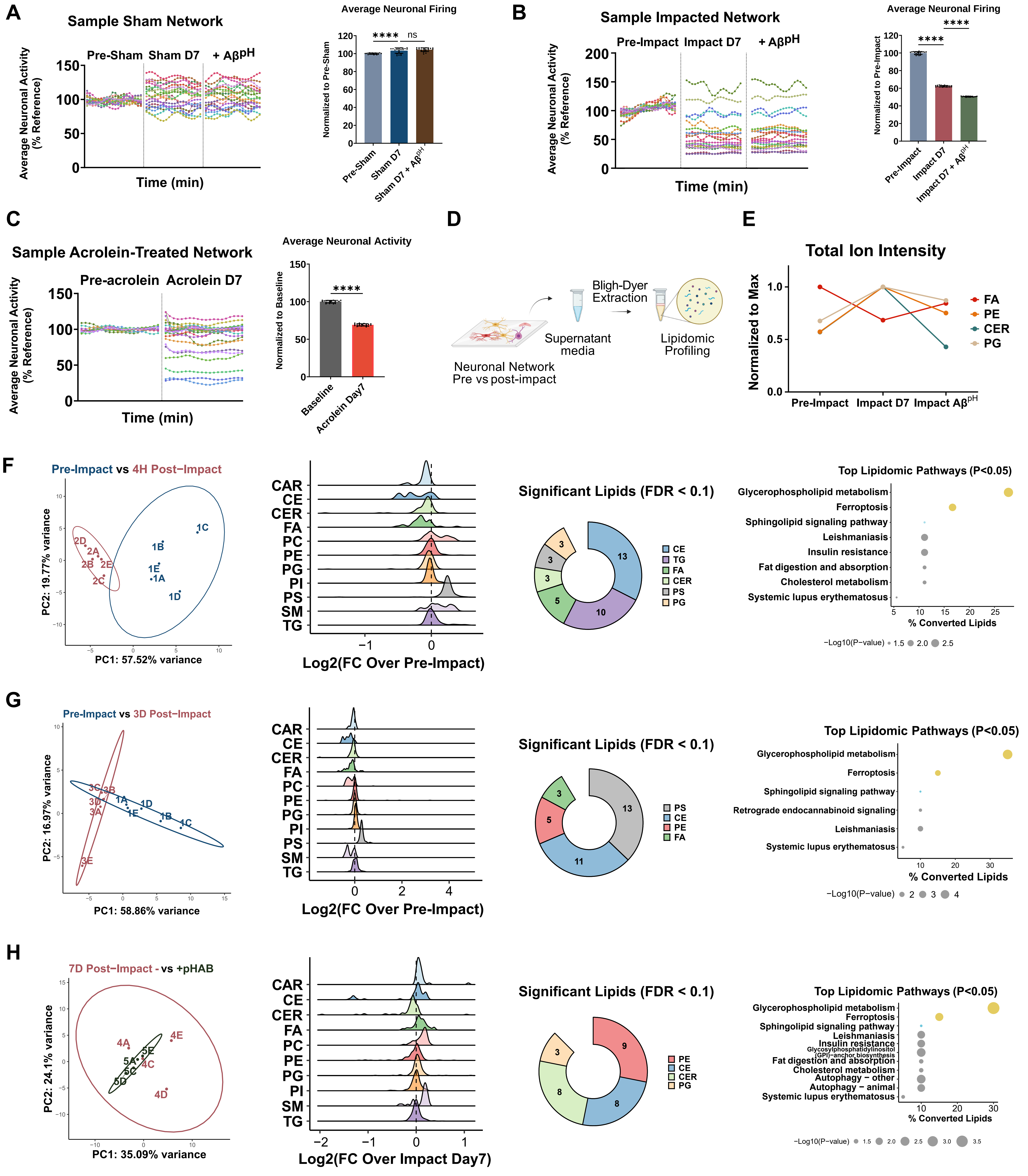
